# Multi-omics Analyses of Facial Skin in Acne Identify Distinct Microbial and Metabolic Features at Lesional and Non-lesional Sites

**DOI:** 10.64898/2026.02.10.702622

**Authors:** Yang Chen, Britta De Pessemier, Tyler Myers, Simone Zuffa, Jasmine Zemlin, Sayeh Pourhamidi, Susi Elaine Dal Belo, Anthony Woo, Magali Moreau, Jolanta Idkowiak-Baldys, Iveta Kalcheva, Paulo Wender Portal Gomes, Chloe Lieng, Samia Almoughrabie, Anna Dan Nguyen, Josh L. Espinoza, Christopher Dupont, Tom Van de Wiele, Chris Callewaert, Daniel McDonald, Karsten Zengler, Andrew Bartko, Luc Aguilar, Philippe Barbarat, Richard L. Gallo, Pieter C. Dorrestein, Qian Zheng, Amina Bouslimani, Se Jin Song, Rob Knight

## Abstract

The microbial and biochemical landscape of clinically normal-appearing skin in individuals with acne remains uncharacterized. Here, we performed longitudinal multi-omics profiling of facial skin from 10 individuals with moderate acne and 10 healthy controls, integrating 16S rRNA gene sequencing, shotgun metagenomics, and untargeted metabolomics across lesional and non-lesional sites. Compositional tensor factorization revealed that non-lesional acne skin occupies a distinct intermediate state between healthy and lesional skin in both the microbiome and the metabolome. Machine learning models distinguished healthy from non-lesional acne skin with 70% accuracy, demonstrating that molecular dysbiosis occurs in areas of the skin without visible lesions. Non-lesional sites exhibited reduced microbial diversity, strain-level shifts in *Corynebacterium* and *Lawsonella* correlating with disease severity, and metabolic alterations, including elevated lipids and perturbed amino acid and dipeptide profiles. Microbe-metabolite co-occurrence network analyses revealed that healthy skin is enriched for protective metabolites such as urocanic acid, while acne-associated skin shows distinct co-occurrence patterns. These findings establish that acne represents a field effect disorder, with molecular alterations extending beyond visible lesions to encompass the entire facial skin ecosystem. This molecular signature of pre-lesional skin provides potential biomarkers for early intervention and suggests that effective acne treatment may require holistic approaches targeting the broader skin environment rather than individual lesions alone.

## MAIN

The skin microbiome supports epithelial barrier integrity and interacts with the cutaneous immune system, thereby influencing skin health and disease. Acne vulgaris is one of the most common skin disorders affecting up to 85% of adolescents and young adults, with potential for lasting personal and social consequences.^1,2^ Despite its prevalence and impact, acne pathogenesis remains incompletely understood and reflects complex interactions between host genetics, hormones, and environmental factors that shape the skin microbiome and trigger inflammation.^3–6^ Microbial shifts, particularly alterations in *Cutibacterium acnes* (*C. acnes*) and *Staphylococcus epidermidis* (*S. epidermidis*) populations, have been documented at lesional sites.^7,8^ These microbial changes can promote inflammation through multiple mechanisms, including activation of innate immune receptors, disruption of the skin barrier, and production of pro-inflammatory compounds.^4^

However, most acne microbiome studies focus exclusively on visible facial lesions, overlooking the broader microbial landscape of non-lesional skin in acne patients. Detailed microbial and biochemical characterization of non-lesional skin has the potential to identify earlier diagnostic biomarkers. Similar to other inflammatory conditions, such as atopic dermatitis (AD)^9^ and inflammatory bowel disease (IBD)^10–12^, individuals with facial acne are expected to have microbial and biochemical changes in the skin across the entire affected area, not just at sites of active inflammation. Thus, we hypothesized that non-lesional skin in individuals with acne, though visually healthy and non-inflammatory, harbors microbial and metabolic alterations reflecting underlying disease processes, changes that precede visible lesion formation. However, these microbial and metabolic signatures remain largely uncharacterized in the context of facial acne. Identifying such changes supports improved understanding of the trajectory of acne development and progress towards novel targets for future microbiome-based preventive interventions.

To address this gap, we employed a multi-omics approach combining microbiome profiling with UPLC-MS/MS untargeted metabolomics. We used two independent 16S rRNA gene sequencing strategies targeting both V1-V3 and V4 hypervariable regions to more comprehensively capture skin-associated taxa. The V1-V3 region provides greater resolution for *Cutibacterium* and *Staphylococcus*,^13^ whereas the V4 region, adopted by the Earth Microbiome Project^14^, offers broader community coverage. We extended this analysis with shotgun metagenomics on a subset of samples to achieve strain-level resolution. We integrated these data layers using dimensionality reduction, machine learning, and multi-omics co-occurrence and network analyses to cross-validate findings and identify associations between microbial community structure and metabolic profiles.

Metabolomic profiling complements microbiome analyses by characterizing the biochemical landscape underlying potential microbial-host interactions. For example, in gastrointestinal disorders such as IBD and Crohn’s disease, untargeted metabolomics has identified candidate diagnostic metabolites and disrupted lipid and amino acid metabolic pathways.^10–12^ The integration of metabolomic data with microbiome composition has proven particularly valuable in understanding how altered microbial communities influence host metabolism, including changes in short-chain fatty acid production, bile acid transformation, and tryptophan metabolism, all of which play critical roles in intestinal inflammation and barrier function.

Similarly, in a study of the microbiome and metabolome of AD, a chronic inflammatory skin condition frequently associated with *Staphylococcus aureus* colonization, distinct metabolic profiles were associated with skin disease states.^9^ Notably, the co-occurrence of bacteria and metabolites was significantly higher in the AD non-lesional skin compared to lesional skin, revealing that apparently healthy skin in AD patients harbors a distinct biochemical landscape with active microbe-metabolite interactions reflective of subclinical disease processes. This finding suggests that metabolic alterations precede or accompany the maintenance of non-inflammatory states in AD-susceptible skin. In AD-lesional skin, *S. aureus* was strongly associated with the presence of dipeptide derivatives, which may contribute to inflammation and barrier dysfunction by modulating immune responses. In contrast, healthy and AD non-lesional skin are enriched for phytosphingosine-derived compounds that correlate with commensal taxa, suggesting that these commensal organisms may actively contribute to skin homeostasis through beneficial metabolite production.^9^

These studies, which leverage microbiome and metabolome data, have been valuable for identifying novel associations between microbial communities and their biochemical outputs; however, such approaches have not yet been applied to facial acne with greater temporal resolution or with systematic consideration of both non-lesional and lesional skin. Here, we conducted a longitudinal study of 10 individuals with clinically active acne and 10 healthy controls over 4 weeks, sampling lesional and non-lesional facial skin sites at multiple time points. Unlike previous cross-sectional studies, our repeated-measures design aimed to capture greater tracking of microbial and metabolic progression across individuals and over time. To account for inter-individual variability and the temporal structure inherent in longitudinal sampling, we applied Compositional Tensor Factorization (CTF),^15^ a dimensionality reduction method specifically designed for repeated-measures data. Our analysis revealed that non-lesional skin has a unique microbial and biochemical profile that can be measurably distinct from that of lesional and healthy skin. Importantly, these findings encourage even more comprehensive spatial and temporal profiling of microbes and their direct relationships with biochemicals, which, complemented by mechanistic studies, can support the implementation of precision or personalized treatments on the skin.

## RESULTS

### Non-lesional acne skin exhibits microbiome and metabolome profiles between healthy and lesional states

To investigate microbial and metabolic differences across acne skin states, we profiled facial skin from 10 women with moderate acne (Global Acne Evaluation score 3) and 10 healthy controls over four weeks. Individuals with acne were sampled from both lesional (AL; papule presence) and non-lesional (ANL; visually healthy-appearing) sites three times weekly, while healthy controls (H) were sampled weekly (**Fig. 1a,b**). H and AL samples were collected from the facial cheeks (left and right), with non-lesional areas representing healthy-appearing skin adjacent to lesional skin in acne individuals. Visual progression of acne lesions were also assigned a severity grading at sampling from 1 (least severe) to 6 (most severe) by trained dermatologists (**Suppl. Fig. 1**). We employed 16S rRNA gene sequencing (independently through V1-V3 and V4 regions), shotgun metagenomics, and untargeted UPLC-MS/MS metabolomics to characterize the microbial and biochemical profile from study participants (**Fig. 1c**).

**Fig. 1.**
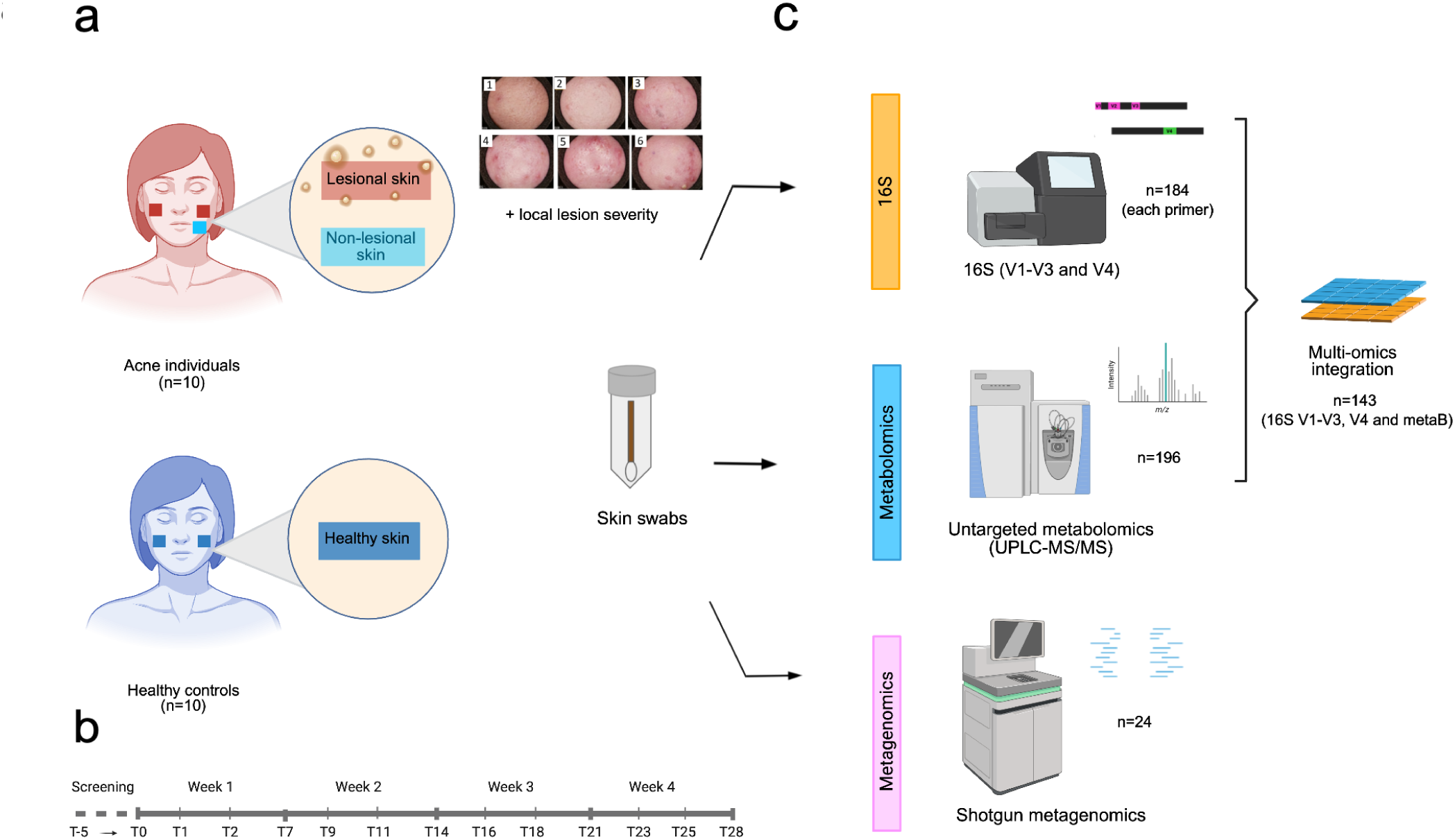
Study design and data characteristics. **(a)** Skin swabs were analyzed from individuals with acne (n=10) and healthy controls (n=10). Acne Lesional (AL) and Acne Non-Lesional (ANL) skin sites were sampled from individuals with moderate acne (GEA 3), and a local severity score (0-6, with 6 being the most severe) was assigned to each swab site. **(b)** Skin samples were collected over four weeks: acne patients were sampled three times per week (a total of 13 time points), and healthy subjects were sampled once per week (a total of 5 time points). 8 time points were selected for AL sites, 5 time points for ANL sites patients, and 3 time points for H subjects were selected for analysis. **(c)** Skin microbiome composition was analyzed using 16S rRNA gene sequencing (V1-V3 and V4 regions), while functional and metabolic profiles were characterized using untargeted metabolomics (UPLC-MS/MS) and shotgun metagenomics. Multi-omics analyses assessed co-occurrences between 16S features and metabolites. *Created with BioRender. De Pessemier, B. (2026)* https://BioRender.com/7yscz20.

First, we used compositional Tensor Factorization (CTF)^15^, a dimensionality reduction method designed for repeated-measures data. Unlike standard ordination approaches that treat each sample as independent, CTF models microbial and metabolic profiles structured within individuals, enabling detection of progressive changes over time and across disease states. To control for potential facial site-specific variation, we analyzed left- and right-cheek samples separately. Across all three data modalities, 16S rRNA gene sequencing (V1-V3 and V4 regions) and untargeted metabolomics, we observed significant separation between H and AL samples at both anatomical sites (**Fig. 2a-f**). Using V1-V3 data, H versus AL separation was significant for both left cheek (*FDR-corrected p = 0.027, F = 3.207*) and right cheek (*p = 0.012, F = 5.303*) (**Fig. 2a,d**). V4 data similarly showed significant H versus AL separation in the left (*p = 0.003, F = 5.182*) and right (*p = 0.012, F = 4.212*) cheeks (**Fig. 2c,f**). Metabolomic profiles distinguished H from AL in both left cheek (*p = 0.027, F = 4.589*) and right cheek (*p = 0.042, F = 3.674*) (**Fig. 2b,e**). Notably, across all three data modalities, acne non-lesional (ANL) samples exhibited a reproducible intermediate clustering pattern between H and AL groups (**Fig. 2a-f**). This pattern achieved statistical significance in the V4 dataset from the left cheek, where ANL samples were significantly differentiated from H samples (*p = 0.022, F = 3.639*) (**Fig. 2c**).

**Fig. 2.**
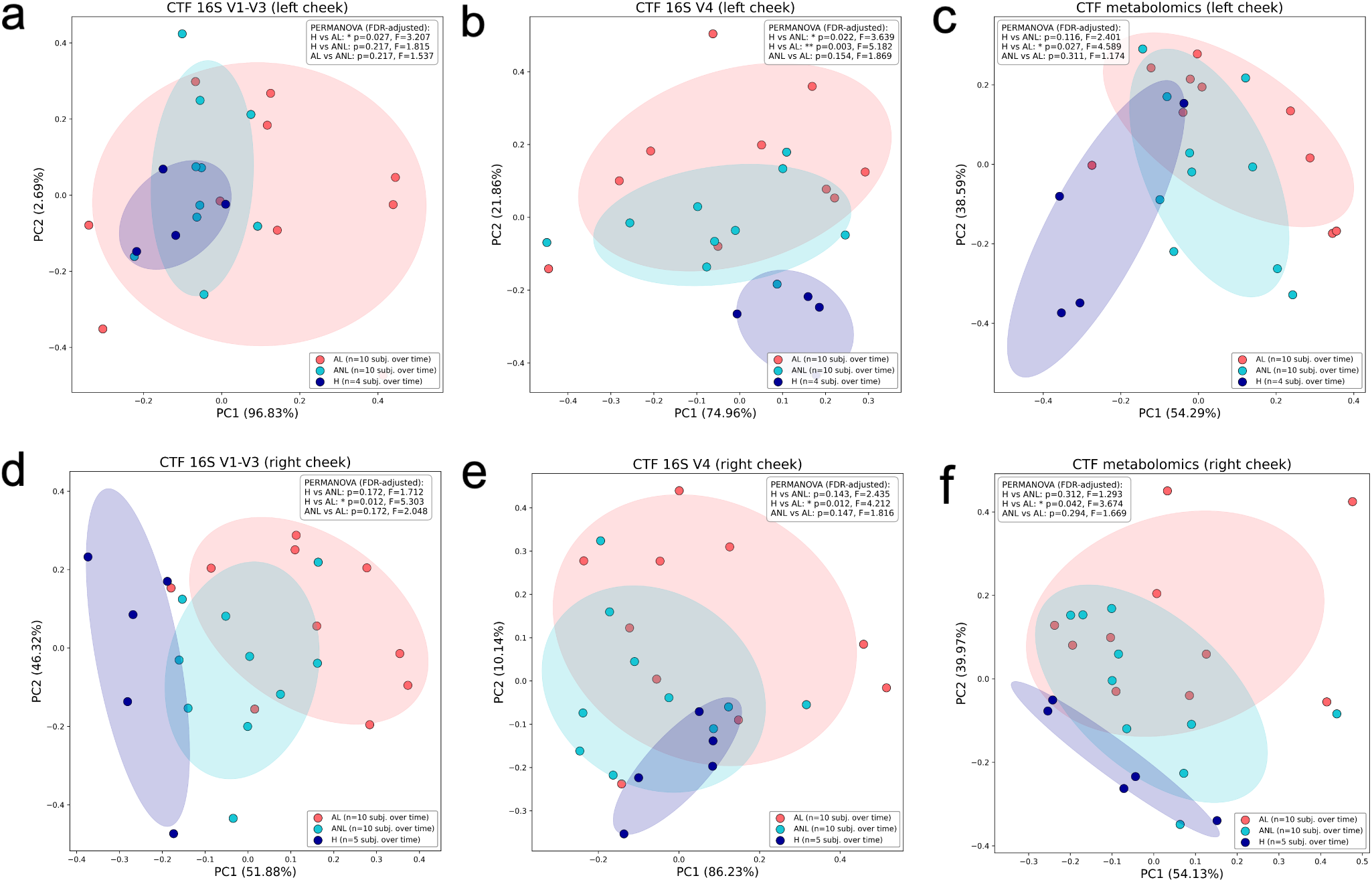
Compositional tensor factorization (CTF) reveals intermediate microbiome and metabolome profiles in non-lesional acne skin relative to healthy and lesional sites. Principal component analysis plots showing compositional tensor factorization (CTF) of **(a-c)** left cheek and **(d-f)** right cheek samples showing **(a,d)** 16S V1-V3, **(b,e)** 16S V4, and **(c,f)** untargeted metabolomics profiles. Samples: H (dark blue), ANL (light blue), and AL (red). Ellipses show 95% confidence intervals. Variance explained is shown on the axes. Group differences assessed by PERMANOVA (F-statistics and FDR-corrected p-values displayed). ***p < 0.001, **p < 0.01, *p < 0.05.

We next examined the ASV-level features contributing most strongly to the CTF model, quantified as the mean loading magnitude across the first three principal component axes for 16S V1-V3 and V4 datasets (**Suppl. Fig. 2a-d**). These loadings reflect taxa that contribute most strongly to overall variance in the CTF models rather than specific disease state associations. Across both primer sets and anatomical sites, the top features included common skin-associated genera (*Staphylococcus*, *Cutibacterium*, *Corynebacterium*, *Streptococcus*, *Lactobacillus*).

### Acne-associated skin displays reduced microbial diversity with strain-level and temporal changes

ANL and AL skin exhibited reduced alpha diversity compared to healthy skin, with this pattern consistent across primer sets and diversity metrics (**Fig. 3a,b; Suppl Fig. 3a,b**). Notably, ANL skin often displayed diversity levels comparable to or even lower than AL skin. In the V1-V3 dataset, Shannon diversity was significantly lower in ANL compared to AL (*p = 0.02*; **Suppl Fig. 3a**), while Faith’s phylogenetic diversity showed significant reductions in both ANL (*p = 0.0175*) and AL (*p = 0.00074*) relative to healthy skin (**Fig. 3a**). The V4 dataset corroborated these trends, with both Shannon diversity (ANL: *p = 0.019*; AL: *p = 0.027*) and Faith’s PD showing consistent reductions in acne-affected individuals (**Fig. 3b; Suppl Fig. 3b**). All statistical analyses employed linear mixed-effects models to account for repeated measurements within individuals.

**Fig. 3.**
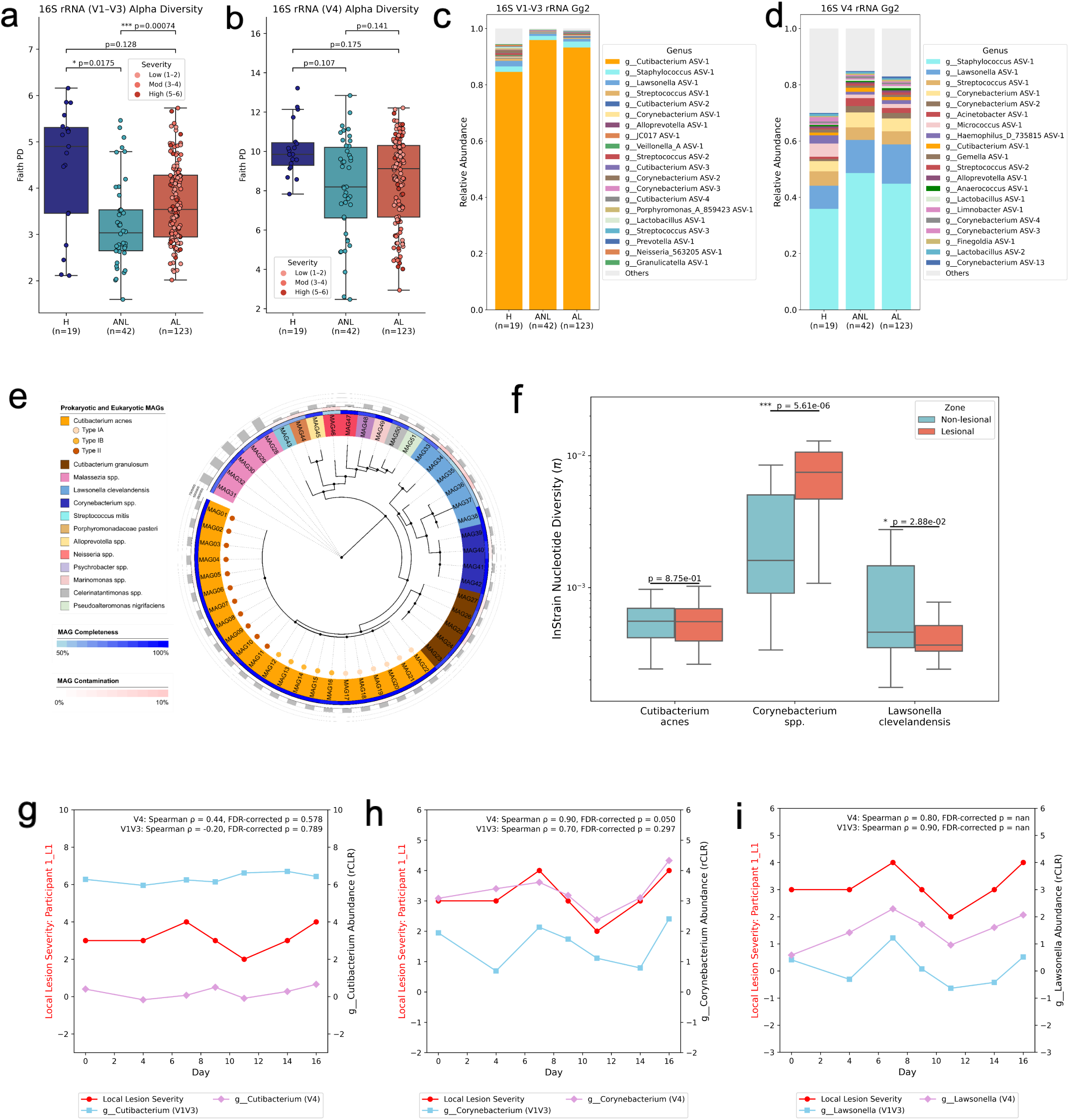
Microbial diversity, composition, and strain variation in healthy and acne-affected skin. **(a-b)** Alpha diversity (Faith’s Phylogenetic Diversity) measured by 16S rRNA gene sequencing using **(a)** V1-V3 and **(b)** V4 primer sets across skin status (H, ANL, AL) and acne severity (Low: 1-2, Moderate: 3-4, High: 5-6). Group differences were assessed using linear mixed-effects models with subject as a random effect; pairwise p-values are shown **(c-d)**. Relative abundance of dominant genera in **(c)** V1-V3 and **(d)** V4 datasets classified from Greengenes2. AL skin shows increased *Staphylococcus* and decreased *Cutibacterium* relative to healthy and non-lesional skin. **(e)** Phylogenetic tree of metagenome-assembled genomes (MAGs) colored by taxon; outer rings indicate completeness and contamination. *Corynebacterium* and *Cutibacterium* species exhibit high diversity and abundance. **(f)** InStrain nucleotide diversity (π) of key skin microbes. *Corynebacterium* spp. and *Lawsonella clevelandensis* show significantly higher strain-level diversity in AL vs. ANL skin (*Mann-Whitney U: Corynebacterium **p = 5.61e-6; Lawsonella p = 2.88e-2*), suggesting strain adaptation in inflamed environments. *C. acnes* shows no significant difference (*p = 0.875*). **(g-i)** Longitudinal tracking of Participant 1 at lesional site L1 (days 0-16) reveals varying relationships between microbial abundance and acne severity.

To characterize strain-level variation in acne-affected skin, we performed shotgun metagenomic sequencing on 24 samples from four acne patients, from lesional and non-lesional sites, reconstructed metagenome-assembled genomes (MAGs), which were taxonomically classified and visualized phylogenetically (**Fig. 3e, Suppl. Fig. 4**). Multi-locus sequence typing (MLST) of *C. acnes* MAGs using the PubMLST scheme^16,17^, which encompasses core metabolic, stress response, and virulence-associated genes, showed Type II strains as most prevalent (n=11), followed by Types IA (n=6) and IB (n=5) (**Suppl. Fig. 5a,b**). While *C. acnes* remained the dominant taxon in both AL and ANL zones (**Suppl. Fig. 5c**), strain-resolved abundance profiles revealed greater variation between individuals than between AL and ANL sites within the same individual (**Suppl. Fig. 5d**). *Cutibacterium*, *Corynebacterium*, and *Lawsonella* species emerged as the most diverse and complete MAGs recovered across subjects (**Fig. 3e).**

We quantified microbial strain variation by calculating gene-level nucleotide diversity (π) using inStrain^18^ for species-level MAGs resolved (**Suppl. Fig. 5e)**. From this analysis, we focused on *C. acnes*, *Corynebacterium* spp., and *Lawsonella clevelandensis*, prioritizing these taxa due to their prevalence, high-quality MAG recovery, and relevance to the skin microbiome. *Corynebacterium* π was significantly elevated in AL compared to ANL skin (*Wilcoxon rank-sum test, p = 5.61e-06*), indicating greater strain heterogeneity at lesional sites, while *L. clevelandensis* π significantly decreased in AL vs. ANL (*p = 0.0288*), suggesting potential strain-level selection or clonal expansion during lesion formation (**Fig. 3f, Suppl. Fig. 5e**). In contrast, *C. acnes* showed no significant differences in π between AL and ANL sites (*p = 0.88*).

To investigate temporal dynamics between microbial composition and disease activity, we performed longitudinal analysis of individuals, examining associations between local lesion severity scores and genus-level 16S microbial relative abundances over time. *Corynebacterium* abundance (RCLR-transformed) significantly correlated with lesion severity in the V4 dataset (*r = 0.90, FDR-corrected p = 0.050*) (**Fig. 3h**; participants 2-4 in **Suppl. Fig. 6**). *Lawsonella* abundance showed a strong positive trend with severity in V4 (*r = 0.80*) (**Fig. 3i**). *Cutibacterium* abundance correlated least strongly with lesion severity (**Fig. 3g**). These observations suggest that the abundances of certain commensal genera can be observed to fluctuate in parallel with local disease activity, whereas *Cutibacterium* abundance remains more stable despite its prevalence in acne-affected skin.

### 16S V4 sequencing achieves the highest classification accuracy for acne detection with strong cross-platform concordance with *Cutibacterium* detected by V1-V3

To evaluate the discriminatory power of each profiling method, we trained random forest classifiers to distinguish between skin phenotypes using 16S V1-V3, 16S V4, and metabolomic features from matched samples. The 16S V4 region consistently outperformed other data types across comparisons, achieving the highest accuracy for separating healthy controls from acne lesional skin *(AUC = 0.67 ± 0.02)* compared to V1-V3 *(AUC = 0.46 ± 0.07)* and metabolomics *(AUC = 0.63 ± 0.02)* **(Fig. 4a)**. V4 similarly showed superior performance in distinguishing healthy from acne non-lesional skin *(AUC = 0.70 ± 0.10)* relative to V1-V3 *(AUC = 0.58 ± 0.12)* and metabolomics *(AUC = 0.44 ± 0.18)*. Notably, classification accuracy was, on average, lowest for the AL versus ANL comparison across all data types, consistent with the subtle biological differences between these spatially adjacent states within the same individuals. Despite this limited classification performance, microbial feature importance rankings were significantly correlated between the V1-V3 and V4 datasets across all three comparisons *(H vs. AL: ρ = 0.552, p = 1.27e-03; H vs. ANL: ρ = 0.439, p = 1.35e-02; AL vs. ANL: ρ = 0.775, p = 3.08e-07;* **Suppl. Fig. 8**), indicating that both primer sets identify consistent acne-associated taxa despite differences in overall classification accuracy.

**Fig. 4.**
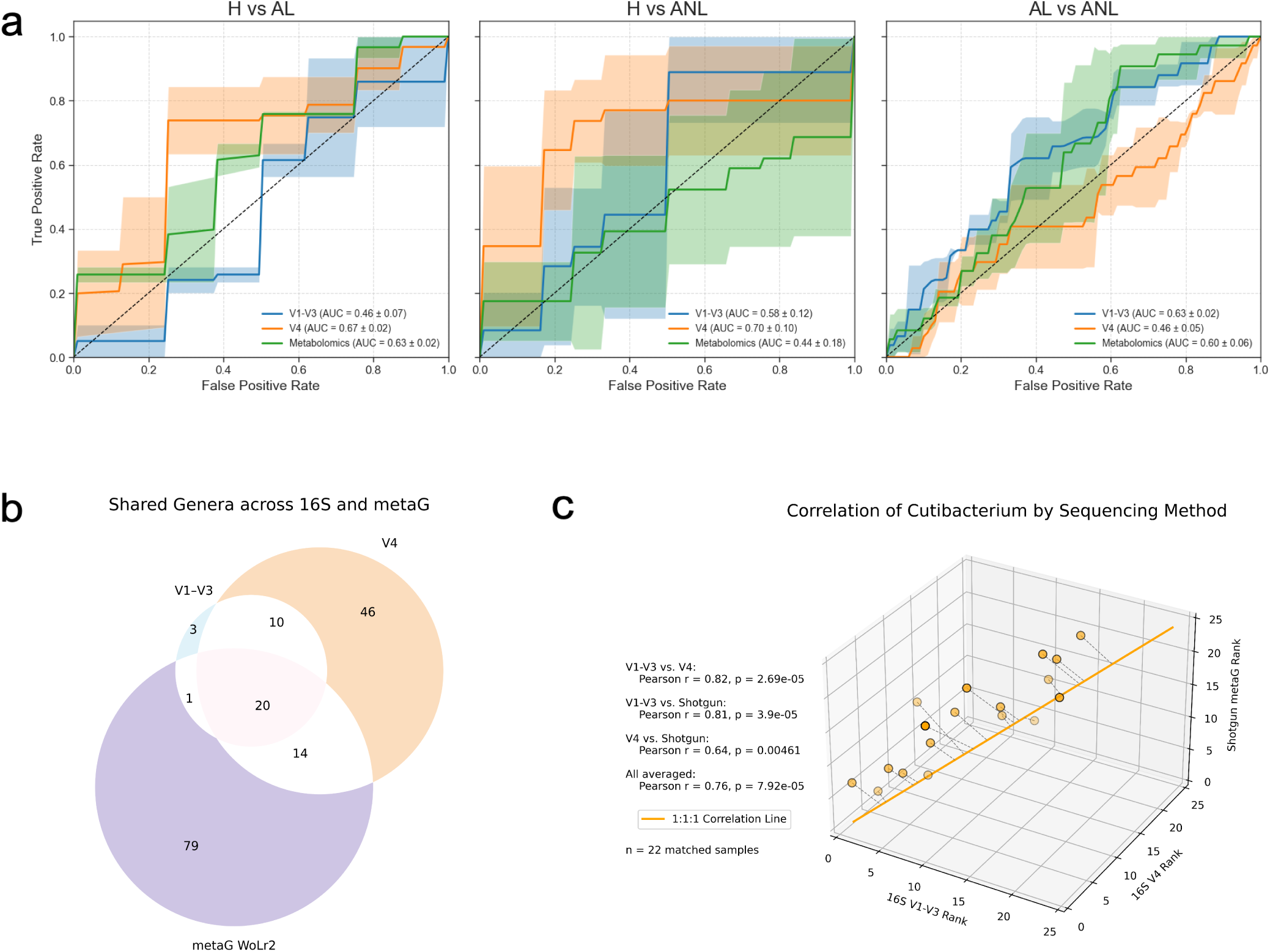
Cross-platform concordance and comparative classification performance of microbiome and metabolomic data in acne phenotyping. Random forest classifications of **(a)** H vs. AL, H vs. ANL, AL vs. ANL. Shaded regions represent 95% confidence intervals. V1-V3 and metabolomics data showed higher predictive performance for H vs. AL and H vs. ANL, while all data types performed similarly in distinguishing AL from ANL. The area under the curve (AUC) values are indicated in the legend. **(b)** Venn diagram showing overlap of detected bacterial genera across 16S rRNA gene sequencing (V1-V3 and V4 regions) and shotgun metagenomic sequencing (WoLr2). **(c)** Concordance of *Cutibacterium* per-sample abundance relative abundance ranks derived from 16S V1-V3, 16S V4, and shotgun metagenomic sequencing across matched samples (n=22).

To compare microbial profiling approaches, we analyzed matched samples collected from four individuals with acne (n = 22; 11 AL, 11 ANL) from 16S rRNA V1-V3, V4, and shotgun metagenomics. Taxonomic classification using the Web of Life 2 reference (WoLr2) database identified 24 genera shared across all platforms, with the V4 region detecting the greatest total number of ASVs (**Fig. 4b**). Three genera were unique to V1-V3, 46 to V4, and 79 to shotgun metagenomics. Direct comparison between primer sets revealed that 16S V4 identified more microbial taxa than V1-V3 (**Suppl. Fig. 7a-b**) and of greater phylogenetic breadth (**Suppl. Fig. 7c-d**). While V1-V3 resolved higher relative abundance of *Cutibacterium* than V4 as previously reported^13^, per-sample ranked abundances of *Cutibacterium* were strongly correlated between the two regions (*r = 0.89, p = 7.87e^−^*^67^; **Suppl. Fig. 7e**), with three-way concordance including shotgun metagenomics also high (*Pearson r = 0.76, p = 7.92e^−^*^05^; **Fig. 4c**). This indicated that although absolute abundances varied between methods, the relative ordering of *Cutibacterium* abundance across samples remained consistent across sequencing platforms.

### Acne skin exhibits elevated sebum and dysregulated lipid and amino acid metabolism

Untargeted metabolomic profiling detected 7,683 consensus MS/MS spectra. Of these, 432 (5.6%) metabolites were confidently annotated through spectral matching to the GNPS2 Standard Spectral Library^19,20^. Inclusion of GNPS Suspect Library nearest-neighbor matches expanded the set of putatively annotated metabolites to 1,032 unique features (13.4%). Chemical class information was further inferred for additional features using SIRIUS^21^ classification using ClassyFire^22^ ontology, resulting in a total of 5,625 total predictions from SIRIUS. Of these, 2,276 SIRIUS predictions which had probability confidence of >75% were retained for analyses (29.6%) (**Suppl. Fig. 9b-c**). Annotations and predictions revealed predominant representation of carboxylic acids and derivatives, amino acids and peptides, fatty acids and derivatives, and alcohols and polyols, consistent with their known roles in skin physiology and barrier function^23,24^ (**Suppl. Fig. 9b-c**).

Partial least squares discriminant analysis (PLS-DA)^25^ of metabolomic features confirmed the clustering patterns observed with CTF (**Suppl. Fig. 10a-b**). Classification performance was strong for H versus AL (*AUC = 0.86±0.03*) and moderate for ANL versus H (*AUC = 0.72±0.05*), but poor for ANL versus AL (*AUC = 0.45±0.03*), consistent with ANL skin being metabolically closer to lesional than to healthy skin (**Suppl. Fig. 10a,b**). Untargeted metabolomics revealed significant biochemical shifts across multiple metabolite classes, with ANL skin exhibiting a unique metabolic profile distinct from healthy controls and AL sites (**Fig. 5**).

**Fig 5.**
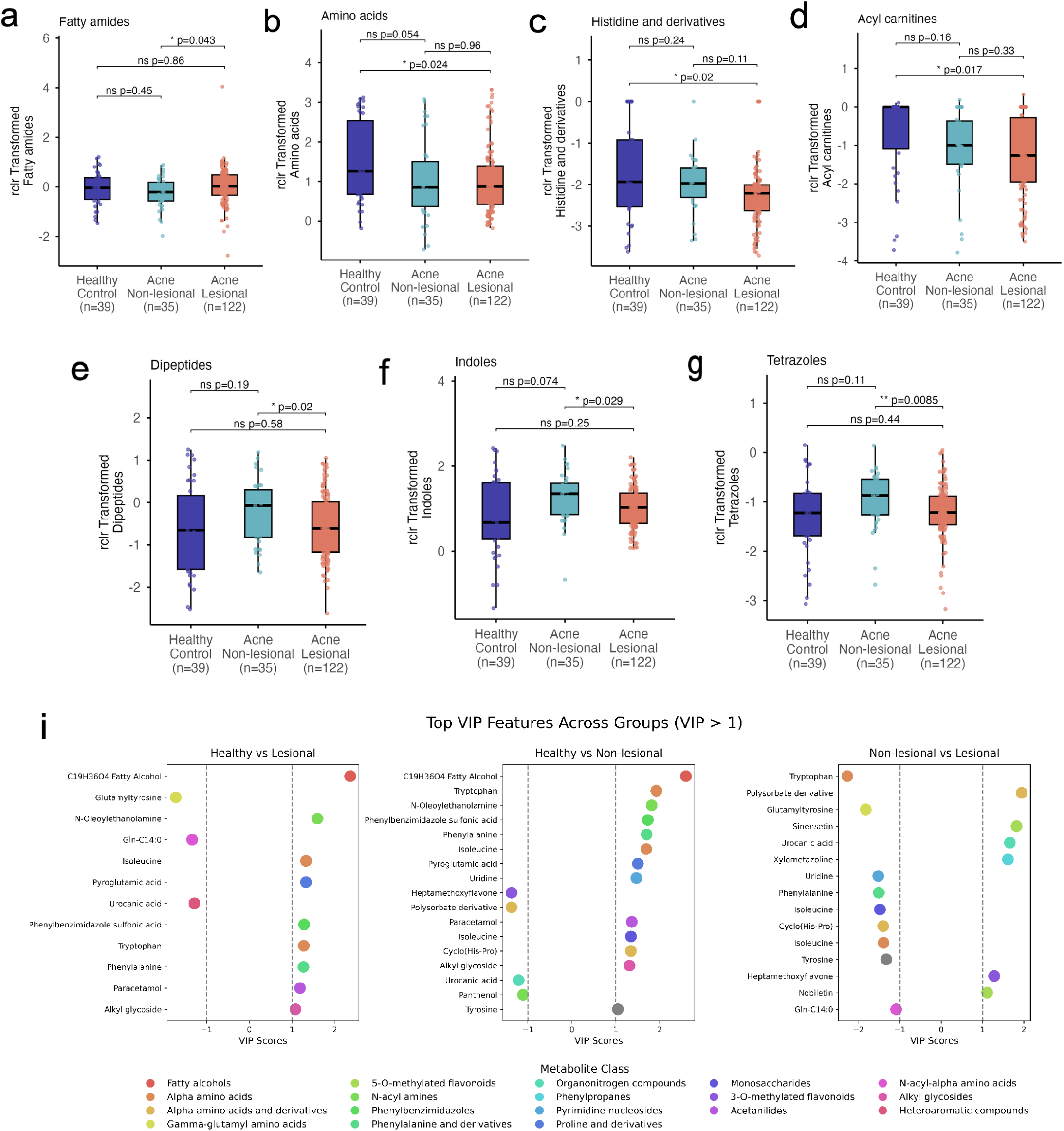
Metabolite class-level and feature-level differences across healthy, acne non-lesional, and acne lesional skin. **(a–g)** Boxplots showing rclr-transformed abundances of major metabolite classes detected by untargeted LC–MS/MS across healthy control (n = 39), acne non-lesional (n = 35), and acne lesional (n = 122) skin. Classes include fatty amides **(a)**, amino acids **(b)**, histidine and derivatives **(c)**, acyl-carnitines **(d)**, dipeptides **(e,g)**, and indoles **(f)**. Points represent individual samples; boxes indicate median and interquartile range. Pairwise comparisons between sites were assessed using linear mixed-effects models (LMMs) with subject ID as a random intercept. P-values were derived from estimated marginal means and adjusted for multiple testing using the Benjamini-Hochberg (BH) method; adjusted *p*-values are shown (ns, not significant). **(i)** Partial least squares–discriminant analysis (PLS-DA) variable importance in projection (VIP) scores for top discriminating metabolites (VIP ≥ 1) across pairwise group comparisons (H vs. AL, H vs. ANL, and ANL vs. AL). Points represent individual metabolite features colored by ClassyFire chemical class.

Fatty amides were significantly elevated in AL compared to ANL (p = 0.043; Fig. 5a), indicating metabolic shifts associated with active inflammation. Clinical measurements confirmed significantly elevated sebum levels (µg/cm²) on the foreheads of individuals with acne compared to healthy controls (*Wilcoxon p = 8.32e-6, ANOVA p = 5.09e-7, F = 26.5*; **Suppl. Fig. 14**), consistent with previous reports^26,27^. AL samples also showed significant reductions relative to healthy controls in amino acids (*p = 0.024*; **Fig. 5b**), histidine and derivatives (*p = 0.02*; **Fig. 5c**), and a trend toward reduced acyl carnitines (*p = 0.017*; **Fig. 5d**). Strikingly, several metabolite classes were significantly elevated in ANL compared to AL skin, suggesting active biochemical processes in clinically normal-appearing skin prior to lesion formation. Dipeptides were significantly higher in ANL than AL (*p = 0.02*; **Fig. 5e**), as were indoles (*p = 0.029*; **Fig. 5f**) and tetrazoles (*p = 0.0085*; **Fig. 5g**).

Untargeted metabolomics revealed significant biochemical shifts across multiple metabolites at class level, with acne non-lesional (ANL) skin exhibiting a unique metabolic profile distinct from both healthy controls and acne lesional (AL) sites (**Fig. 5**). Strikingly, several metabolite classes were significantly elevated in ANL compared to AL skin, suggesting active biochemical processes in clinically normal-appearing skin prior to lesion formation. Dipeptides were significantly higher in ANL than AL (*p = 0.02*), as were indoles (*p = 0.029*) and tetrazoles (*p = 0.0085*). Conversely, fatty amides were significantly elevated in AL compared to ANL (*p = 0.043*), indicating metabolic shifts associated with active inflammation. AL samples also showed significant reductions relative to healthy controls in amino acids (*p = 0.024*), histidine and derivatives (*p = 0.02*), and a trend toward reduced acyl carnitines (*p = 0.017*). These findings suggest that ANL harbors a distinct biochemical landscape characterized by heightened metabolic activity.

We next extracted features from the PLSDA and examined metabolites with variable importance in projection (VIP) scores ≥1 (n = 24). While the previous analysis examined metabolite classes, this feature-level analysis revealed that individual discriminative metabolites also fell predominantly into lipid and amino acid-related categories. The classes of the metabolites with VIP scores ≥1 from the PLS-DA model were predicted via ClassyFire^22^ (**Fig. 5i**). In the H versus AL comparison, key discriminative metabolites included fatty alcohols (C₁₉H₃₆O₄), N-acyl amines (N-oleoylethanolamine), and amino acids and amino-acid derivatives (glutamyltyrosine, (iso)leucine, pyroglutamic acid, tryptophan, phenylalanine), with phenylbenzimidazole sulfonic acid and urocanic acid also among top features (**Fig. 5i**, left panel). The H versus ANL comparison revealed a similar pattern, with fatty alcohols (C₁₉H₃₆O₄), tryptophan (an amino acid with tryptophan group), and N-oleoylethanolamine, phenylbenzimidazoles (phenylbenzimidazole sulfonic acid, phenylalanine), and amino acids ((iso)leucine, pyroglutamic acid, uridine) as top discriminative features, alongside cyclo(His-Pro), paracetamol, and alkyl glycosides (**Fig. 5i**, middle panel). The ANL versus AL comparison showed the weakest differentiation, with tryptophan, polysorbate derivative, glutamyltyrosine, sinensetin, urocanic acid, and phenylalanine as top features (**Fig. 5i**, right panel). The consistent appearance of lipid-related metabolites (fatty alcohols, N-acyl amines) and amino acids across both H vs. ANL, H vs. AL, and ANL vs. AL comparisons demonstrates that metabolic alterations characteristic of acne lesions are detectable at the molecular level in visually healthy skin.

### Significant metabolite features driving acne phenotype separation and their multi-omics co-occurrence patterns with skin microbes

We then applied MMVEC (Microbe-Metabolite Vectors)^28^, a neural network method that models conditional co-occurrence probabilities, to the 24 high-VIP metabolites and 10 frequently top-ranked microbial taxa across pairwise comparisons (H vs. AL, H vs. ANL, ANL vs. AL; **Fig. 6, Suppl. Fig. 11**). To investigate metabolite origins, we performed domain-specific MASST^29,30^ searches against publicly available datasets, yielding matches across microbial monocultures (microbeMASST)^31^, microbial communities (microbiomeMASST), human/murine tissues (tissueMASST)^32^, personal care products (PCP), plants^33^, plant sources (plantMASST) ^33^, and food sources (foodMASST)^34^. MASST results were subsequently overlaid onto the MMVEC-derived heatmap to highlight potential microbial sources of these metabolites.

**Fig. 6.**
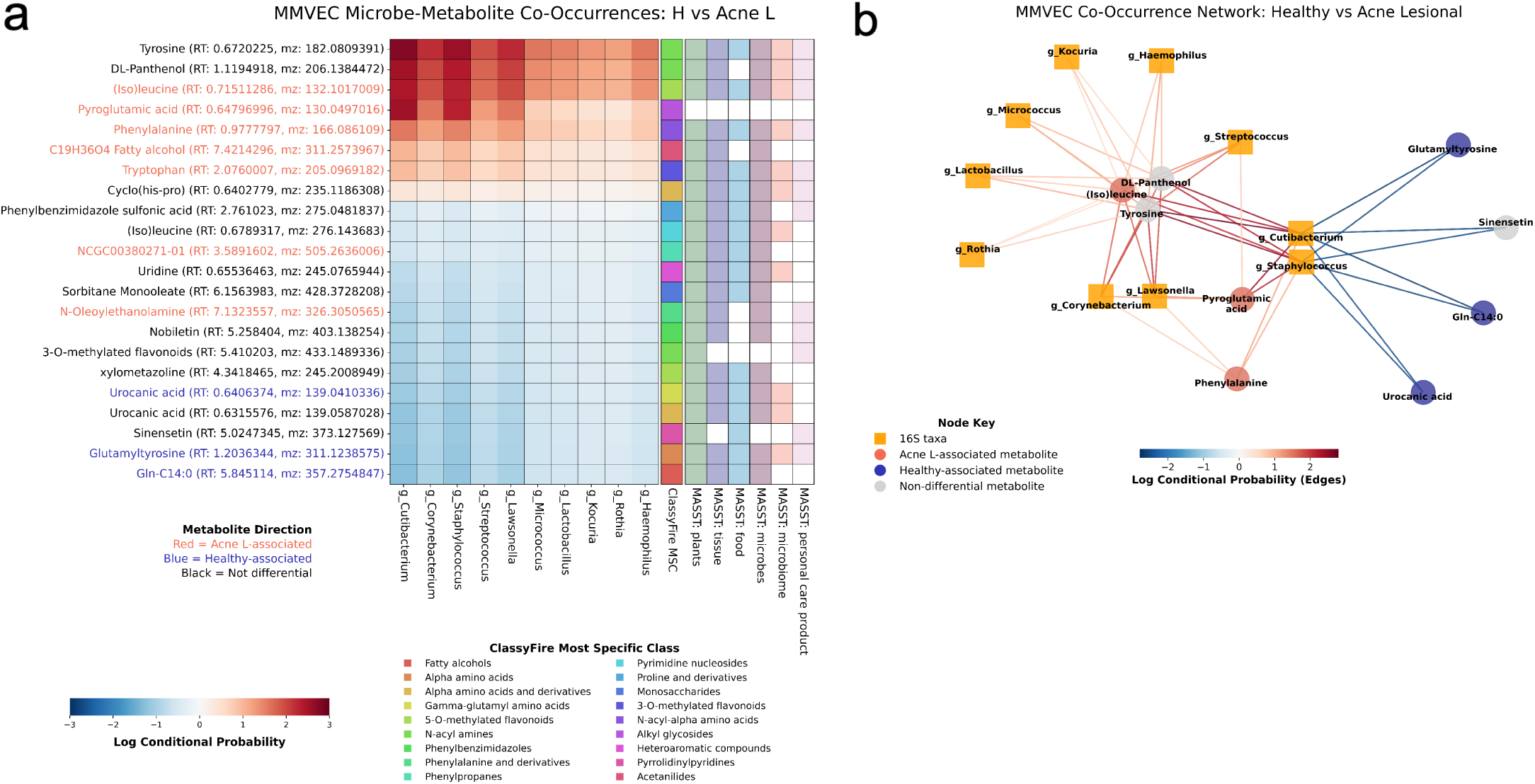
Microbe-metabolite co-occurrences of acne lesional vs. healthy skin. **(a)** Heatmap of log conditional probabilities for co-occurrence between the top 30 differentially abundant metabolites and 10 key taxa. Metabolites are annotated by identity, retention time (RT), and m/z, with color coding indicating VIP directionality (red=AL-associated, blue=H-associated, black=non-differential). Right annotation bars show ClassyFire classifications and MASST database matches. **(b)** Network of top 20% microbe-metabolite associations comparing H vs. AL skin. Taxa (orange squares) and metabolites (circles) are colored by VIP association: red (AL), blue (H), gray (non-differential). Edge weight and color represent log conditional probability of co-occurrence, with red indicating AL-enriched and blue indicating healthy-enriched associations.

Amino acids elevated in acne, tyrosine, phenylalanine, (iso)leucine, and tryptophan, showed positive co-occurrences with all ten genera. MASST confirmed their presence in skin-associated bacteria: tyrosine in *Propionibacterium*, *Micrococcus*, *Staphylococcus*, *Lactobacillus*, and *Streptococcus*; phenylalanine in *C. acnes*, *S. epidermidis*, *Streptococcus*, and *Lactobacillus*; (iso)leucine in *Lactobacillus* spp.; and tryptophan in *C. acnes*, *S. epidermidis*, *Corynebacterium*, *Lactobacillus*, and *Streptococcus*. These amino acids were also detected in plants, mammalian tissues, food, and personal care products, suggesting potential multifactorial origins. Importantly, we note that MicrobeMASST spectral matches do not establish *in situ* microbial production on the skin, as many reference spectra derive from *in vitro* microbial cultures and may reflect media components or shared background metabolites rather than active microbial biosynthesis *in vivo*.

N-oleoylethanolamine, elevated in both AL and ANL skin, showed negative co-occurrence with all microbes and was detected across human skin, microbial cultures, plants, and personal care products. Notably, this compound has been identified in skin with primary microbial infections and in samples from patients with inflammatory bowel disease^35^, rheumatoid arthritis, and osteoarthritis^35,36^, suggesting it may serve as a broader inflammation marker. The dipeptide cyclo(His-Pro), enriched in ANL versus H skin and showing positive co-occurrence with all microbes, was detected in *Propionibacterium*, *S. aureus*, *Micrococcus*, and *Streptococcus* monocultures, as well as in skin affected by primary infections, indicating potential value as an early inflammatory marker.

Metabolites elevated in healthy skin, urocanic acid, glutamine-C14:0, and glutamyltyrosine, showed negative co-occurrence with all bacteria. Urocanic acid was identified across food, microbes, plants, and human tissue, though microbial hits did not overlap with genera in our analysis. Glutamine-C14:0, detected in microbes, tissue, plants, and food, can be produced by *Staphylococcus* monocultures despite showing negative co-occurrence in our data. Glutamyltyrosine was identified in human skin via tissueMASST.

Personal care product (PCP)-associated metabolites, spanning cosmetics, hair care, oral care, and household products, were observed across multiple individuals with significant inter-individual abundance variation (Kruskal-Wallis p<0.05; **Suppl. Fig. 12**), consistent with prior findings that such compounds persist on skin for weeks despite daily cleansing^37^.

## DISCUSSION

This study reveals that acne-associated microbial and metabolic disruption extends beyond visible lesions to clinically normal-appearing skin. Using integrative multi-omics approaches, we demonstrate that non-lesional skin in acne patients exhibits its own unique microbiome and metabolomic profile, reinforcing the importance of holistic skin microbiome sampling in acne studies.

While the cohort size was modest (10 acne patients and 10 healthy controls), the strength of our analyses lies in its several methodological approaches, which we cross-compare. Additionally, our longitudinal design, with multiple sampling points over four weeks, enabled capture of dynamic changes in individual patients. Critically, we employed CTF that accounts for subject-specific variation, revealing consistent microbial differences that conventional alpha diversity metrics failed to detect. We benchmarked two 16S rRNA primer sets (V1-V3 and V4), demonstrating that V4 offers broader taxonomic resolution and compatibility with large-scale datasets while maintaining strong per-sample correlation with V1-V3 and shotgun data for key taxa like *Cutibacterium*. This dual-primer validation extends the acne-associated microbial signature beyond *Cutibacterium* to include additional taxa of interest.

Untargeted metabolomics revealed significant metabolic features in acne skin, reinforcing the hypothesis that acne-associated metabolic dysbiosis extends beyond visible lesions. Fatty amides were significantly enriched in lesional skin of acne patients, a finding that aligns with their well-documented role in sebum composition and microbial metabolism.^24^ We observed increased levels of N-oleoylethanolamine in both AL and ANL skin compared to healthy skin. N-oleoylethanolamine is an agonist of peroxisome proliferator-activated receptor alpha (PPARα)^38,39^, a receptor known to play a key role in sebocyte differentiation, sebum production, and inflammation, processes central to acne pathogenesis^40,41^. PPARα also regulates lipid metabolism, epidermal differentiation, and cellular processes such as proliferation and migration^40^. N-oleoylethanolamine’s role may be similar to leukotriene B4 (LTB4), another PPARα ligand that promotes inflammation and sebum production, and whose inhibition via the drug zileuton has been shown to significantly improve acne symptoms ^42–44^. We also identified the N-acyl lipid glutamine-C14:0, a molecule not widely studied in dermatology but that may be involved in lipid metabolism and inflammatory processes in acne.^45^

We observed altered levels of amino acids, amino acid derivatives, and dipeptides, suggesting metabolic changes. Tryptophan, an amino acid containing an indole group, was significantly elevated in acne skin. Impaired metabolism of branched-chain amino acids and tryptophan has been implicated in acne pathogenesis, severity, and scarring^46^. Tryptophan and its metabolites are endogenous ligands for the aryl hydrocarbon receptor (AhR), a transcription factor that regulates immune responses and skin barrier function. AhR activation influences T cell differentiation, promoting regulatory T cells or pro-inflammatory Th17 responses depending on the ligand and tissue context. In acne, elevated tryptophan may contribute to inflammation through AhR-mediated immune dysregulation, potentially skewing T cells toward pro-inflammatory phenotypes^47,48^. This suggests that tryptophan metabolism represents a molecular link between microbial activity, host metabolism, and the inflammatory response in acne lesions.

(Iso)leucine levels were elevated in both lesional and non-lesional acne skin compared to healthy controls, consistent with previously reported increases in serum^46^ and with increased isoleucine biosynthesis pathways reported in psoriasis.^5^ These branched-chain amino acids play important roles in dermal collagen synthesis^49^. Phenylalanine was enriched in both lesional and non-lesional acne skin, and tyrosine was significantly elevated in non-lesional skin compared to both healthy and lesional sites. Conversely, urocanic acid levels were higher in healthy skin compared to both acne groups, and higher in lesional skin compared to non-lesional skin. Phenylalanine can be converted to tyrosine via phenylalanine hydroxylase^50^. Together with tryptophan and urocanic acid, these aromatic amino acids participate in UV protection^51–53^. Urocanic acid also plays a role in defense against pathogens, while tryptophan functions as an antioxidant. Phenylalanine and tyrosine serve as precursors of melanin^46^ and a previous study similarly reported phenylalanine upregulation in serum of acne patients^54^.

Our MMVEC analysis revealed positive co-occurrence patterns between specific skin commensals (*Cutibacterium*, *Staphylococcus*, *Micrococcus*) and amino acids enriched in acne-affected skin. MicrobeMASST searches confirmed that these amino acids are associated with skin-resident microbes, suggesting potential microbial contributions to their presence. Additionally, the dipeptide cyclo(His-Pro) was elevated in non-lesional acne skin, indicating its potential as a pre-lesional biomarker. These results align with findings from Gomes *et al.*^9^, who similarly reported elevated dipeptide derivatives in atopic dermatitis, highlighting a broader role for altered dipeptide metabolism in inflammatory skin diseases.

Additionally, several metabolites identified using MASST matched compounds found in personal care products, consistent with previous findings by Bouslimani *et al*.^23,37^, which demonstrated that these products significantly influence the skin’s molecular composition alongside skin cells and microbial communities. Notably, microbial shifts linked to personal care products were more pronounced in occluded areas such as armpits and feet compared to the face^37^. Despite an eight-week wash-out period, these compounds remained detectable, highlighting the need to account for exogenous compounds as potential confounders in future metabolomic studies.

Several important limitations should be considered when interpreting these findings. First, although our longitudinal design captured repeated sampling across lesional, non-lesional, and healthy skin, the study was not designed to track site-specific conversion from non-lesional to lesional skin. The observed microbial and metabolic differences in clinically non-lesional skin therefore reflect phenotypic associations with acne status at the time of sampling rather than predictive biomarkers of lesion formation. Second, the acne cohort was selected to enrich for individuals exhibiting dynamic changes in lesion severity, which may amplify observed contrasts and limit generalizability to individuals with stable or mild disease. Third, while CTF accounts for repeated measures at the subject level, several downstream analyses necessarily pooled longitudinal samples and therefore did not fully model within-subject correlation. Fourth, shotgun metagenomic analyses were limited to a subset of individuals, thereby constraining statistical power to detect subtle or taxon-specific effects. Fifth, the study population comprised young adult women from a single geographic region, limiting extrapolation across sex, age, ethnicity, and environmental contexts. Sixth, untargeted metabolomic features reflect a complex mixture of host-derived, microbially associated, and environmentally sourced compounds, and the persistence of personal care product metabolites underscores the challenge of disentangling endogenous skin biology from exogenous exposures. Finally, the observational nature of this study precludes causal inference regarding whether microbial or metabolic alterations initiate, amplify, or result from acne-associated inflammation.

Despite these limitations, our findings advance the field by applying microbiome-metabolome integration to facial acne with temporal resolution and by systematically considering both non-lesional and lesional skin, an approach not previously undertaken. Unlike previous cross-sectional studies, our repeated-measures design enabled tracking of microbial and metabolic dynamics across individuals and over time. The application of CTF, specifically designed for repeated-measures data, enabled us to account for inter-individual variability and temporal structure. Importantly, our demonstration that non-lesional skin possesses a unique microbial and biochemical profile, measurably distinct from both lesional and healthy skin, underscores the need for comprehensive spatial and temporal profiling of microbes and their direct relationships with biochemicals. Future studies should incorporate larger and more diverse cohorts, site-matched tracking of lesion emergence, and mixed-effects statistical frameworks to validate candidate biomarkers. Complemented by mechanistic and interventional studies, such integrated multi-omics approaches can clarify causal relationships between the skin microbiome, metabolism, and acne pathophysiology, ultimately supporting the development of precision and personalized treatments for acne.

## METHODS

### Study Design

This study was conducted by L’Oreal in Bucharest, Romania, between January and March 2022, following approval from the IBL Life Ethics Committee (EC21-COS-070) and in accordance with the Declaration of Helsinki and Good Clinical Practice guidelines. Informed consent was obtained from all participants prior to enrollment.

Thirty-one women aged 18-24 years were recruited at the Centre International de Développement Pharmaceutique (CIDP) in Bucharest, including 16 participants with clinically diagnosed moderate facial acne (GEA 3) and 15 healthy controls without clinical signs of acne (not matched to acne participants). A dermatologist confirmed all acne diagnoses. The local severity of lesion sites where swabs were collected was quantified using a clinician-assigned score ranging from 0 (absent) to 6 (highest severity) (Suppl. Fig. 1). Non-lesional skin was defined as healthy-appearing skin located at least 5 cm from any visible lesion. Standardized full-face and close-up lesion photographs documented clinical presentation and local lesion severity.

Participants maintained their usual hygiene routines but avoided additional skincare products and makeup on sampling days, using only a gentle face wash the day before each visit. Exclusion criteria included current use of topical acne treatments, oral acne medications (e.g., isotretinoin), or antibiotics. Sebum levels (µg/cm²) were measured on each participant’s forehead at every time point using a Sebumeter.

### Sample Collection and Selection

All subjects were sampled three times per week for 4 weeks. For the acne group, samples were collected from three facial zones: two lesional sites (papule-containing areas on the left and right cheeks) and one non-lesional site (healthy-appearing skin on the cheek). Healthy controls were sampled from two facial zones (left and right cheeks).

Due to budgetary constraints, we selected a subset of participants for sequencing and LC-MS/MS analysis from the original cohort of 16 acne patients and 15 healthy controls. We selected 10 acne patients who demonstrated the greatest change in lesion severity over time and 10 healthy controls selected at random. For acne patients, we used a two-step selection process: first, we identified subjects with a change ≥2 in at least one lesional area, yielding 11 eligible participants. We then refined our selection using combined criteria, including subjects if either the sum of lesion severity scores from both lesional areas (C1+C2) exceeded 35 or the sum of all control area scores (C3) was ≤1. This process resulted in our final sample of 10 acne patients.

The final sample set included eight time points from lesional sites and five time points from non-lesional sites for each of the 10 acne patients, plus three time points from each of the 10 healthy controls. At each sampling site and time point, we collected dual swabs: one for microbiome analyses (16S rRNA gene sequencing and shotgun metagenomics) and one for metabolomics.

All skin sampling was conducted in a climate-controlled room maintained at 21 ± 1°C and 50% relative humidity. For microbiome analysis, samples were collected using a sterile dual-swab pre-moistened with 95% ethanol solution. Each swab was rubbed firmly on the designated site for 60 seconds, covering a 2 cm² surface area. Following collection, each cotton swab was placed into a cryotube, immediately flash-frozen in liquid nitrogen, and stored at −80°C. Samples were subsequently shipped to the Center for Microbiome Innovation (CMI) at the University of California San Diego for downstream processing and analysis.

### 16S rRNA Gene Sequencing and Data Analysis

DNA was extracted from swabs using the standard MagMAX Microbiome Ultra Nucleic Acid Isolation Kit (ThermoFisher Scientific, Waltham, MA, USA) on the KingFisher Flex instrument (ThermoFisher Scientific, Waltham, MA, USA), as previously described^55^. Serially diluted Lysobacter sp. OAE881 with known cell counts was used as a positive control in the KatharoSeq protocol, which aids in setting a limit of detection for low-biomass samples^55^, for both 16S V1-V3 and 16S V4 runs.

The 16S rRNA V1-V3 and V4 hypervariable regions were amplified using the following primer sets using the Human Microbiome Project^56^ protocol for V1-V3 and the Earth Microbiome Project^57^ protocol for V4 (515f-806r with Golay barcodes): V1-V3 Region Forward primer AGAGTTTGATCCTGGCTCAG, Reverse primer TTACCGCGGCTGCTGGCA, and V4 Region Forward primer GTGYCAGCMGCCGCGGTAA, Reverse primer GGACTACNVGGGTWTCTAA. Amplicon libraries were equal volume pooled, PCR cleaned using the QIAquick PCR Purification Kit (QIAGEN, Hilden, Germany), and quantified using the Qubit dsDNA Quantification Assay Kit (ThermoFisher Scientific, Waltham, MA, USA). Amplicons were then sequenced on an Illumina MiSeq platform at the UCSD Sequencing Core using a 300-cycle reagent kit (v2) (Illumina, San Diego, CA, USA). V1-V3 and V4 datasets were generated from independent sequencing runs.

Raw sequencing data for both the V1-V3 and V4 amplicons were processed using the Qiita platform (study ID 14901) with its standard QIIME 1-based 16S rRNA gene analysis workflow Sequences were quality filtered using a PHRED score cutoff of ≥ 4 to retain as much data as possible^58^, then trimmed to 150 bp before denoising. The Deblur algorithm was applied, resulting in a feature table of amplicon sequence variants (ASVs)^59^. Reads classified as ‘Mitochondria’ or ‘Chloroplast’ were removed from the dataset. Due to the expected low biomass in these samples, we employed the KatharoSeq protocol^55^. This process utilizes a set of positive control dilutions to assess the minimum number of reads a sample should contain. This threshold was calculated to be 11,054 reads for the V1-V3 data and 3,769 for the V4 data. Taxonomic classification was then performed using the Greengenes2^60^ 2022.10 full-length backbone Naive Bayes classifier, with phylogenetic placement inferred independently via SEPP^61^ fragment insertion into the Greengenes2 full-length 16S rRNA gene backbone through q2-fragment-insertion.

### 16S Microbiome Analyses

16S microbiome analyses were first conducted to investigate differences in the microbial communities between samples from H subjects, ANL, and AL skin groups from both V1-V3 and V4 datasets. To increase statistical power and account for temporal sampling, all samples collected across time points were aggregated by skin status group (H, ANL, AL), resulting in a high number of samples per group for cross-sectional comparisons.

Alpha diversity was assessed using Faith’s Phylogenetic Diversity^62^, a phylogenetically informed metric that leverages the feature table and Newick tree from Qiita^63^ with Greengenes2^60^ 2024.9 as well as the Shannon index^64^, a metric that measures richness and evenness. Pairwise statistical significance was assessed using the Mann-Whitney U-test^64^. *p<0.05* was considered statistically significant, with all p-values labeled on the plots. Multiple samples from each individual were included in this analysis to examine the large-scale effects of acne on skin microbial diversity. Linear mixed-effects models with participant-specific random intercepts were used to account for repeated sampling, and pairwise group comparisons were assessed via Wald tests corrected with the Holm-Bonferroni method. p<0.05 was considered statistically significant, with all p-values labeled on the plots.

We assigned taxonomy ASV features within V1-V3 and V4 datasets at Kingdom, Phylum, Class, Order, Family, and Genus levels. Each table was then collapsed based on assigned taxonomy labels. The Genus-level collapsed table was used to visualize the relative abundance of microbes with stacked bar plots. The top 15 genera detected with each primer set were highlighted, with matching genera between V1-V3 and V4 datasets assigned the same color. All remaining genera were grouped under the “Others” category.

### Shotgun Metagenomic Sequencing

To assess whether microbial differences between AL and ANL skin were more pronounced at the species or strain level, we performed shotgun metagenomic sequencing on a subset of samples. We selected 24 samples (12 AL, 12 ANL) from the four acne patients who showed the largest cumulative change in lesion scores over the study period. For each patient, we sampled one lesional and one non-lesional site at three specific time points: the lowest lesion score before peak, the peak lesion score, and the lowest score after peak. This sampling strategy was aimed at capturing full trajectories of lesion development and resolution.

DNA libraries were prepared from samples using 1 ng input DNA following the Nextera XT library preparation kit protocol (Illumina, cat. no. FC-131-1096) and sequenced on an Illumina MiSeq platform with a paired-end 150 V2 kit (Illumina, San Diego, CA, USA). All 48 paired-end samples were sequenced to a minimum depth of 15 million reads. Per-sample paired-end raw sequencing fastqs were quality-assessed using FastQC^65^ (v0.12.1) with default parameters and summarized with MultiQC^66^ (v1.23). All fastqs passed quality control metrics for per-base sequence quality and per-sequence quality scores. Raw reads were then human DNA-filtered using Method 1 from Guccione et al. (2025)^67^, which involves sequential alignment to GRCh38 and T2T-CHM13 using minimap2, followed by indexing-based filtration with the Human Pangenome Reference^67,71^ (HPRC) using movi^72^. After removal of all human reads, microbial reads were retained only when achieving ≥10% genome breadth using micov^73^ to minimize multimapping-driven artifacts.

### Metagenome Assembly

Metagenome assembly and binning were performed using the Viral Eukaryotic microbial Archaeal (VEBA)^74,75^ (v2.2.0) open-source software suite. Quality-filtered reads were assembled using VEBA’s assembly module, which implements metaSPAdes from SPAdes^76,77^ (v3.15.5) with default parameters (k-mer sizes: 21, 33, 55, 77, 99, 127). Metagenome-assembled genomes (MAGs) were recovered from assembled contigs with VEBA’s binning module which uses MaxBin2^78^ (v2.2.7) and MetaBAT2^79^ (v2.15). MAG quality was assessed using CheckM2^80^ (v1.0.1) for prokaryotic genomes, CheckV^81^ (v1.0.1) for viral genomes, and BUSCO^82^ (v5.4.3) for eukaryotic genomes to maximize recovery of high-quality MAGs across all domains of life.

Following assembly and binning, all MAGs were filtered to keep those with at least 50% completion and no more than 10% contamination. 22 *C. acnes* MAGs were typed using the Multi-Locus Sequence Typing (MLST) scheme^16^ for *C. acnes* using the *C. acnes* PubMLST^16,17^ database. This web browser tool allows for the categorization of an assembled *C. acnes* genome file into one of the known *C. acnes* strain types using *aroE, aptD, gmk, guaA, lepA, recA, sodA, tly*, and *camp* marker genes.

### LC-MS/MS Metabolomics Sample and Data Processing

The cotton buds of all samples (57 H, 50 ANL, and 159 AL; a total of 266 samples) were added into a 96-deep well plate (2 mL). Using a multichannel pipette, 500 μL aliquots of Methanol (MeOH)/H_2_O (1:1) were added to each well for extraction. The plates were sonicated for 10 min in an ultrasound bath, Branson 2800 (Danbury, CT, USA), vortexed (∼10 s), and then the swabs were removed with tweezers, and the extracts were dried using a LabConco Centrivap. The dried extracts were stored at −80 °C until resuspension. The samples were resuspended with 100 μL of MeOH/H_2_O (1:1) spiked with 1 mM of sulfadimethoxine (internal standard), then the plates were sonicated for 5 min in an ultrasound bath, Branson 2800 (Danbury, CT, USA), vortexed (∼10 s), and centrifuged for 20 min at 2,000 rpm and 4 °C. Using a multichannel pipette, 100 μL of each sample was transferred to a new Eppendorf 96-shallow well microplate for LC-MS/MS analysis. A mix of MeOH/H_2_O (1:1) containing 1 mM of sulfadimethoxine was used as a blank, and a mix of sulfamethazine (C12H14N4O2S), sulfamethizole (C9H10N4O2S2), sulfachloropyridazine (C10H9ClN4O2S), sulfadimethoxine (C12H14N4O4S), amitriptyline (C20H23N·HCl), and coumarin-314 (C18H19NO4) was used as quality control (QC). A QC pool containing 5 μL of all samples was also created.

The metabolomic analyses were performed in a Vanquish UHPLC system coupled to a Q-Exactive orbitrap mass spectrometer (Thermo Fisher Scientific, Waltham, MA, USA). The chromatographic analysis was carried out on a Kinetex Polar C18 column (100 × 2.1-mm, 1.7-μm particle size, 100-A pore size, Phenomenex, Torrance, CA, USA). A high-pressure binary system was used for gradient elution. The column and autosampler were kept at 30 °C, and 25 °C, respectively. The flow rate was 0.5 mL/min, and the elution was carried out using ultra-pure water (solvent A) and acetonitrile (solvent B), both acidified with 0.1% formic acid (FA). The gradient method was set as follows: 0-1.0 min, 5% B; 1.0-8.0 min, 5%-99% B; 8.0-9.0 min, 99%-5% B; 9.0-10.0 min, 5% B to stabilize the system before the subsequent analysis.

For mass spectrometry analyses, data-dependent acquisition (DDA) was employed in an *m/z* range of 100 to 1500 Da, using an electrospray source operating in positive ionization mode. Before data acquisition, sodium formate solution (Thermo Fisher Scientific, San Diego, CA, USA) was used for external calibration with an error rate of less than 0.5 ppm. The spray voltage was set to 3.5 kV, the sheath N2 gas flow rate was set to 53 psi, and the auxiliary N2 gas pressure was set to 14 psi. The ionization source was maintained at 270 °C, a 60 V S-lens RF level was applied, and the auxiliary gas heater was set to 440 °C. Full-scan MS1 was performed at 1.0 × 10^6^ with a resolution of 35,000 and a maximum ion injection of 100 ms. MS2 experiments were performed with a resolution of 17,500 and a maximum IT time of 100 ms. TopK6 was used to select the six most abundant precursor ions per MS1. The MS2 precursor isolation window was set to 2 Da with an offset of 0.5 Da. The normalized collision energy was set to a ramp from 20 to 40 eV, and the exclusion (MS1 and MS2) for unassigned ion charge states was set to 5 S-Lens, as well as isotope peaks.

The LC-MS/MS data were converted from RAW standard data format (Thermo Fisher Scientific, Waltham, MA., USA) to mzML format using MSConvert 3.0.2^83^. The LC-MS/MS data were processed using MZmine 3.1.0^84^. The mass detection of MS1 and MS2 levels was performed using a signal noise of 3.0E4 and 2.0E3, respectively. The ADAP chromatogram builder was used to build the chromatogram, and a minimum group size of scans was set to 4, the minimum intensity of the group to 1.0E5, and the highest to 5.0E4 with an *m/z* tolerance of 10 ppm. The ADAP resolver module (wavelets) was applied to chromatographic deconvolution. The local minimum resolver module was used for chromatographic resolving. The chromatographic threshold was set to 90% with a minimum absolute feature height of 6.0 × 10^4^, a minimum peak top/edge ratio of 1.40, a peak duration ranging from 0.00 to 3.0 min, and a minimum scan time of 3. The isotope peak grouper module was applied to detect the isotopes with a *m/z* and RT tolerance of 3.0 ppm and 0.2 min (charge 1 was used as a standard) for the most intense isotope. To remove duplicate features and aligners, a 5.0 ppm tolerance with a 4-minute window was used, and the weights for *m/z* and RT were set to 3:1, respectively. The peak list was exported as a .mgf file and a .csv file containing feature information, which was used for downstream statistical analysis.

The outputs containing MS features were then used in the Feature-Based Molecular Networking (FBMN) workflow on the GNPS2 platform^19,20^. The parameters used were set as follows: the tolerances for the precursor ion mass (MS1) and the MS2 fragment ions were set to 0.02 Da each. A cosine of 0.6 and a minimum of 5 matching peaks were applied to create the molecular network and identify library matches. ^85^. The *in silico* GNPS2 Suspect Library was also used to generate annotations^86,87^. Statistical analyses were performed in R version 4.3.3.

### Metabolomics Feature Filtering and Annotation

We obtained a total of 7,683 unique MS2 spectra and employed multiple state-of-the-art workflows to annotate metabolic features. First, we utilized the GNPS Standard Spectral Library, a database containing MS2 spectra acquired by reference standards. Additionally, we leveraged the *in silico* GNPS Suspect Spectral Library^86^, which was generated via a nearest-neighbor search algorithm, to expand metabolite annotation beyond the standard reference library. To further classify unknown small molecules, we used SIRIUS 4. Specifically, the CANOPUS^88^ module was used to predict compound molecular classes using ClassyFire^22^ ontology. For these predictions, we retained only the ones where the Most Specific Class had a confidence score of at least 75%. This threshold was determined based on visual inspection of the probability distribution across all features within each MostSpecificClass category (Suppl. Fig. 10c) and the exploratory nature of this investigation. Using this approach, we predicted the classes of 2,276 features (29.6%) for downstream analysis. To visualize the chemical diversity of these metabolites, we constructed a Sankey plot displaying their ClassyFire Superclass, Class, and Subclass categories, including only nodes representing at least 1% of the total annotated metabolites. Additionally, we visualized the molecular network using Cytoscape 3.9.1^89^, where metabolites were mapped based on their ion mass size and annotated Superclass category to explore structural and functional relationships.

Using R version 4.3.3, we examined the Total Ion Count (TIC) ordered by sample run sequence across different sample types: (a) Subject samples, (b) Pooled quality control samples, (c) Sixmix quality control samples, (d) Blank samples, and (e) Blank control swabs samples. To ensure data quality, we removed 37 subject samples with TIC values between 6 × 10⁸ and 5 × 10⁹, thereby reducing the dataset from 266 to 229 samples. After sample removal, we assessed the internal standard sulfadimethoxine, yielding a coefficient of variance of 12.6%. To ensure high-confidence metabolite features, we applied additional filtering before statistical analyses. We removed features where the ratio of their mean intensity in pooled quality control samples to blank samples (Pool_Blank=Mean_pool / Mean_blank) was less than 5, pooled samples to sixmix samples (Pool_Mix=Mean_pool / Mean_sixmix) was less than 5, and pooled samples to control samples (Pool_CTL=Mean_pool / Mean_ctl) was less than 5. This step helped eliminate potential contaminants and technical artifacts, refining the dataset for downstream analysis. To further refine the dataset, we removed contaminant polyethylene glycol (PEG) features using both targeted and untargeted homolog discovery approaches. Specifically, we identified PEG homologs with detectHomologues() from the package homologueDiscoverer^90^ using a targeted approach based on known mass differences, followed by an untargeted approach to detect additional homologous series. After applying these quality filtering steps, we retained 1,688 high-confidence metabolite features for downstream analysis. PCA was then used to assess sample-level structure and revealed a subset of samples exhibiting clustering patterns that were not explained by host identity, plate number, or health status, suggesting the presence of unaccounted technical or biological variation. To account for this, 33 samples were removed, resulting in a final dataset of 196 samples. Following this additional filtering step, 1,142 metabolites remained for analysis.

### Metabolomics Analyses

For each pairwise comparison, a Partial Least Squares-Discriminant analysis (PLS-DA) using the plsda() function from the mixOmics^91^ package (R v4.3.3) were conducted on robust center log-ratio (rCLR)-transformed data. PLS-DA was run with four components (ncomp = 4) on a scaled (scale = TRUE) data matrix. The explained variance of each component was assessed, and a variable importance in projection (VIP) score threshold of 1 was applied to the first component to select the most relevant metabolic features.Model performance was evaluated using the perf() function with 5-fold cross-validation (M-fold, 100 repeats) (Suppl. Fig. 10). To assess classification accuracy, the classification error rate (CER), balanced error rate (BER), and area under the curve (AUC) were computed (Suppl. Fig. 10). Scatter plots of the first two components from PLS-DA were generated to visualize class separation, with group centroids representing the mean position of each category.

We then focused on the specific metabolite features from the pairwise PLS-DA model with VIP scores above 1 (including those near the threshold). Because PLS-DA can be prone to overfitting, we also performed univariate statistical tests to confirm their significance in distinguishing among skin groups. Each significant metabolite was annotated according to its ClassyFire category, and a vertical heatmap was generated to display the range of p-values for each feature based on the feature table, including those near the 0.05 cutoff.

To assess differences in metabolite classes and individual metabolite features across AL, ANL, and healthy skin regions while accounting for repeated measurements within participants, linear mixed-effects models were fitted using the lme4 package (v1.1.37)^92^. Skin site was included as a fixed effect, and subject ID was included as a random intercept. Post-hoc pairwise contrasts were performed using the emmeans package (v1.11.1)^93^, with p-values adjusted for multiple testing using the Benjamini–Hochberg procedure.

To determine the potential origin of metabolites detected in the skin metabolome, we performed MASST (Mass Spectrometry Search Tool) batch searches using the domainMASSTs tool (https://github.com/robinschmid/microbe_masst)^31–34^. MS/MS spectra were searched against the latest indexed data (metabolomicspanrepo_index_latest). Searches were conducted using the following parameters: minimum cosine score of 0.7, a precursor *m/z* tolerance of 0.05 Da, a fragment ions *m/z* tolerance of 0.02 Da, and a minimum of 3 matched signals. A maximum of 5 parallel queries were allowed, and previously searched spectra were skipped. Analog searching was disabled.

### Multi-omics Analyses

#### Compositional Tensor Factorization

Given the longitudinal aspect of the 16S and metabolomics data with multiple samples per subject, we applied compositional tensor factorization (CTF)^15^ to account for host-based variation over time and reduce the dimensionality of the data to one representative sample per individual. CTF is specifically designed for compositional microbiome data with repeated measures, decomposing the data tensor into latent factors that capture variation across features, samples, and conditions while preserving the compositional nature of the data.

Using the same feature tables from the 16S rRNA primer comparison and metabolomics analyses, we implemented CTF using the Gemelli package (v0.0.12) with default parameters. To best align with the CTF methodology and account for the paired nature of AL and ANL samples from the same anatomical location, we ran CTF separately for left and right cheek samples. This approach ensures that lesional and non-lesional comparisons are made within the same facial region, controlling for potential site-specific microbiome differences.

For each cheek, CTF constructs a three-dimensional tensor with dimensions corresponding to features (ASVs or metabolites), samples, and conditions (skin groups: H, ANL, AL). The method then performs a robust centered log-ratio (RCLR) transformation to handle the compositional nature of the data, followed by tensor decomposition using alternating least squares to identify latent factors that explain the major patterns of variation.

Samples were aggregated based on host subject ID and skin group (healthy, acne NL, acne L) across all longitudinal time points in the study. This aggregation strategy allows CTF to model within-subject variation while identifying consistent patterns that distinguish between skin conditions. The resulting ordination captures the primary axes of variation while accounting for the repeated sampling design and compositional constraints inherent in microbiome and metabolomics data.

After dimension reduction, we evaluated the clustering of healthy, non-lesional, and lesional samples using permutational multivariate analysis of variance (PERMANOVA) with 999 permutations, as implemented in the scikit-bio^94^ Python package (version 0.5.8). P-values were corrected for multiple comparisons using the false discovery rate (FDR) method to account for testing multiple pairwise comparisons.

#### Microbe-Metabolite Co-occurrences

To investigate potential functional links between microbial taxa and metabolite features, we performed co-occurrence analysis using the MMVEC algorithm^28^ with the paired-omics function, setting latent_dim=3, learning_rate=1e-3, batch_size=50, input_prior=1, summary_interval=1, and -p-epochs=500. We selected the following 10 microbial taxa, *Micrococcus*, *Corynebacterium*, *Cutibacterium*, *Staphylococcus*, *Haemophilus*, *Streptococcus*, *Kocuria*, *Lawsonella*, *Rothia*, and *Lactobacillus*, for focused downstream analysis. These Genera taxa were chosen based on their frequent presence among the top 15 ranked features (in either the V1-V3 or V4 16S rRNA gene datasets) across each of three key comparisons: (1) H vs. AL, (2) ANL vs. ANL, and (3) overall top discriminatory features. Seven of the ten taxa met the top 15 threshold in all three comparisons. In contrast, *Kocuria*, *Rothia*, and *Lactobacillus* met the criterion in two of three cases but were retained due to their consistent signals and established biological relevance to the skin microbiome. We subsequently focused on the metabolites identified in our PLS-DA analyses (VIP scores > 1), integrating them with the MMVEC output. The resulting microbe-metabolite associations were visualized in a heatmap (Fig. 6b), where each cell represents the log conditional probability of a given metabolite conditioned on the presence of a particular microbial genus.

#### Cross-Platform Concordance Analyses

To assess the concordance of microbial community composition across sequencing platforms, we compared genus-level detection and abundance patterns among 16S rRNA gene sequencing (V1-V3 and V4 hypervariable regions) and shotgun metagenomic sequencing. This analysis utilized a subset of matched samples (n=22; 11 AL, 11 ANL) from four individuals with acne for which all three sequencing modalities were available. Shotgun metagenomic reads were taxonomically classified using the Web of Life 2 reference (WoLr2) database.

Genus-level overlap across platforms was visualized using Venn diagrams. To evaluate quantitative concordance, we calculated Pearson correlation coefficients on the ranked relative abundances of individual genera between platforms. For two-way comparisons (e.g., V1-V3 vs. V4), we report the Pearson correlation coefficient (r) of the per-sample ranked relative abundances. For three-way comparisons incorporating shotgun metagenomics data alongside both 16S primer sets, we fit a three-dimensional linear regression model and report the coefficient of determination (*R²*) as a measure of the variance in ranked relative abundances explained across all three platforms.

#### Random Forest Classifications

To assess the ability of microbial and metabolite profiles to differentiate between skin groups, we implemented a Random Forest (RF) classification strategy using scikit-learn^95^ (v0.24.1). This approach was applied independently to genus-level 16S rRNA data from both V1-V3 and V4 regions, as well as to untargeted metabolomics profiles. Feature tables were imported from QIIME 2^96^ (v2024.2.0) artifacts and converted to pandas (v1.5.3) DataFrames, with taxonomy annotations at 16S ASV level. Only samples with complete metadata and skin group labels were retained for analysis.

To prevent overfitting and account for repeated measures from the same individuals, classification was performed using a group-stratified 5-fold cross-validation. In each fold, approximately 80% of the data was used for training and 20% for testing, as determined by setting GroupKFold(n_splits=5). All samples from a given subject (host_subject_id) were assigned entirely to either the training or testing set to prevent information leakage.

Random Forest models were implemented using RandomForestClassifier with 1,000 estimators (n_estimators=1000) and a fixed random seed (random_state = 42) to ensure reproducibility. Each model was trained on the training set for a given fold and used to predict probabilities on the test set. For each binary pairwise comparison (e.g., H vs AL), class labels were binarized (0 vs 1) and Receiver Operating Characteristic (ROC) curves were computed. The Area Under the ROC Curve (AUC-ROC) served as the primary evaluation metric for classification performance. If any fold contained a training set with only one class, that fold was skipped.

To identify features that contributed most to classification, we extracted feature importances based on the mean decrease in Gini impurity. These importance scores were calculated for each fold and averaged across folds to obtain a consensus ranking of features. The top-ranked genera or metabolites were visualized to highlight the most discriminatory features across data modalities.

## ADDITIONAL

### Data and code availability

Raw mass spectrometry data generated in this study will be deposited in the MassIVE repository under an accession ID during publication. The 16S and metagenomic data can be found in Qiita study 14901 and EBI-ENA accession PRJEB106897 ERP187942. The metabolomics GNPS job can be found at the following link https://gnps2.org/status?task=4b0e3f9e93c845ed8103bf7fdd5e7021.

Scripts for all other analyses, as well as the code used to produce manuscript figures, can be found in our GitHub repository: https://github.com/knightlab-analyses/Acne_Longitudinal_Microbiome.

### Disclosures

SP, SDB, AW, LA and PB are employed by the company L’Oréal Research and Innovation, Aulnay-sous-Bois. MM, JIB, QZ and AB are employed by the company L’Oréal Research and Innovation, Clark, NJ. L’Oréal Research and Innovation, as funder, has been involved in the decision to submit the study for publication.

R.K. is a scientific advisory board member, and consultant for BiomeSense, Inc., has equity and receives income. He is a scientific advisory board member and has equity in GenCirq. He has equity in and acts as a consultant for Cybele. He is a Vice President and board member of Microbiota Vault, Inc. He is a board member of N=1 IBS advisory board and receives income. He is a Senior Visiting Fellow of HKUST Jockey Club Institute for Advanced Study. The terms of these arrangements have been reviewed and approved by the University of California, San Diego in accordance with its conflict of interest policies.

R.G. is a co-founder, scientific advisor, consultant, and equity holder in MatriSys Biosciences and is a consultant, receives income, and equity holder in Sente Inc.

P.C.D. is an advisor and holds equity in Cybele, BileOmix, and Sirenas, and a Scientific co-founder, advisor, holds equity, and/or received income from Ometa, Enveda, and Arome with prior approval by UC-San Diego. P.C.D. also consulted for DSM animal health in 2023.

D.M. is a consultant for BiomeSense, Inc., has equity, and receives income. The terms of these arrangements have been reviewed and approved by the University of California, San Diego, in accordance with its conflict-of-interest policies.

C.C. is the founder and receives income from Phiome.

## Acknowledgments

Funding was provided by the Center for Microbiome Innovation and by L’Oréal USA S/D, Inc. B.D.P. was supported by the Research Foundation Flanders (grants 1S04122N and V477223N).

