## Supplementary material for "Multi-omics Analyses of Facial Skin in Acne Identify Distinct Microbial and Metabolic Features at Lesional and Non-lesional Sites": Suppl. Fig. 1-13

### EXTENDED DATA FIGURES

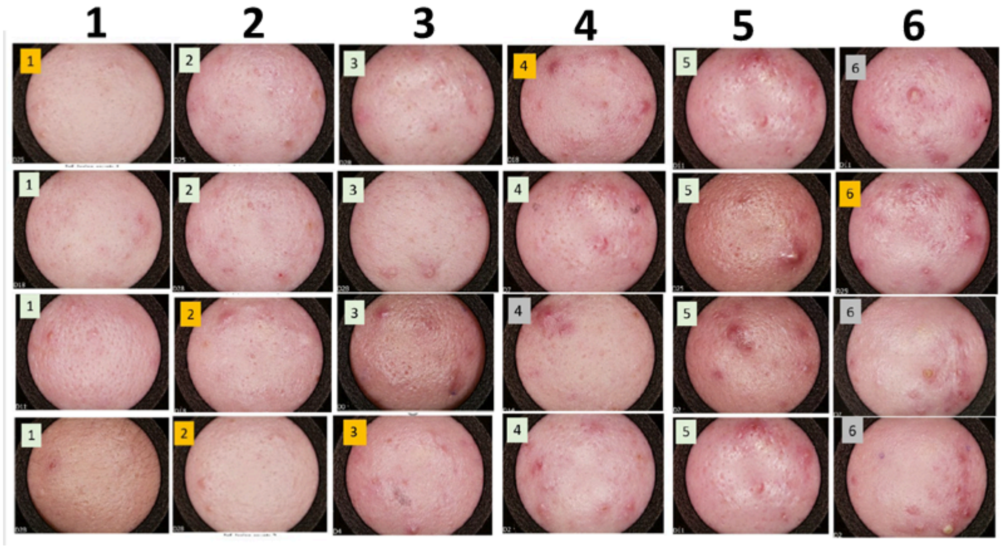


**Suppl. Fig. 1. Visual progression of acne severity across local lesion severity grades.** Representative clinical images of lesional skin from individuals with moderate acne (GEA 3), arranged by local severity grade from 1 (least severe) to 6 (most severe). Each column corresponds to a single severity grade, and each row represents a different subject with a similar lesion severity. Grade 1 shows primarily small comedones and minimal inflammation, while higher grades (e.g., 5 and 6) exhibit nodular lesions, papules, pustules, and greater erythema. The images highlight the morphological variability present within each clinical localized grade that was used for subsequent analyses in this study.

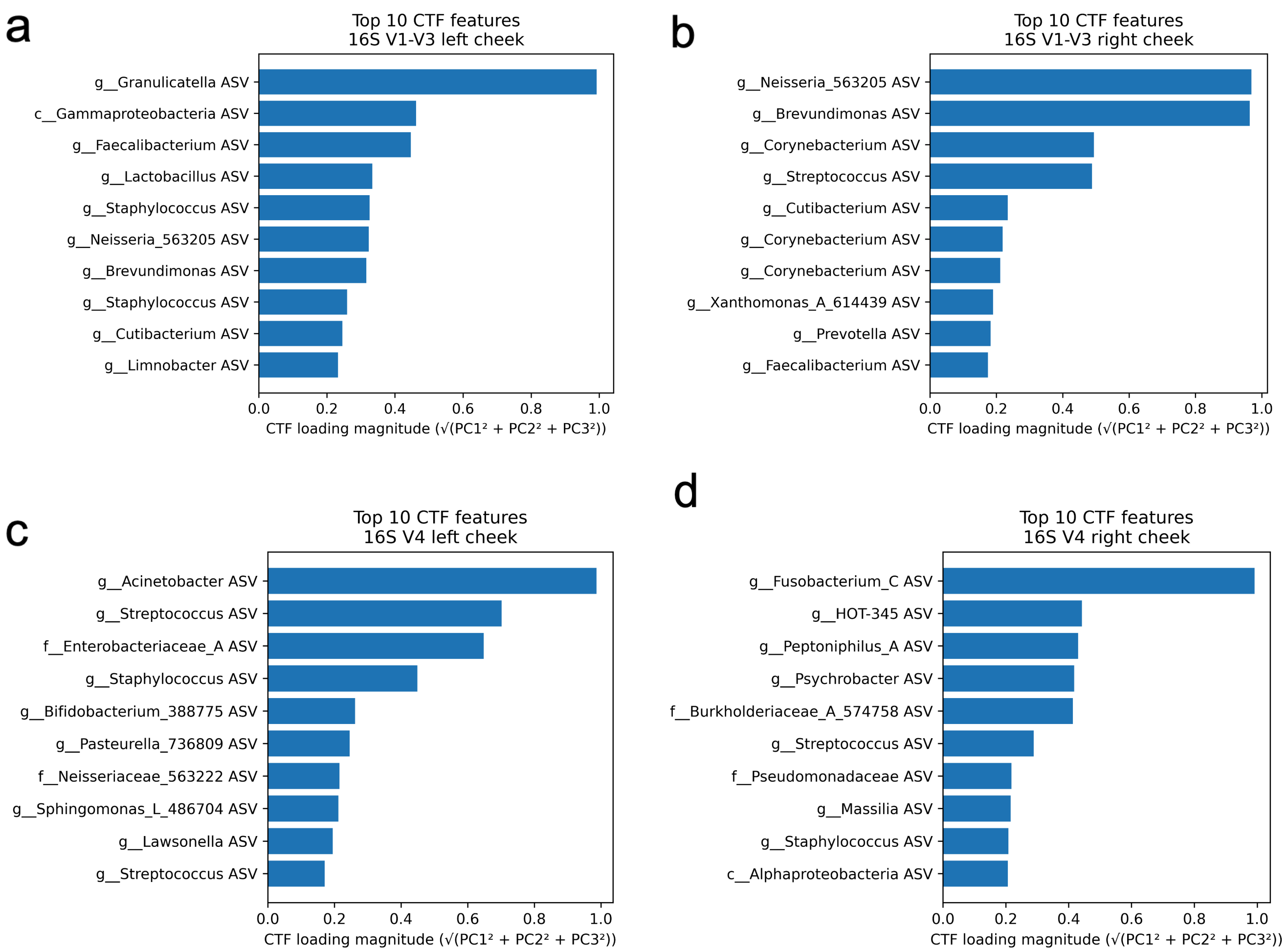


**Suppl. Fig. 2. Top CTF 16S V1-V3 and V4 ASV feature loadings.** Bar plots show the top 10 features with the largest CTF loading magnitudes ($\sqrt{PC1^2 + PC2^2 + PC3^2}$), stratified by 16S rRNA gene primer set and cheek sampling side. Panels **a)** and **b)** show results for the V1-V3 region from the left and right cheek, respectively, while panels **c)** and **d)** show results for the V4 region from the left and right cheek. Features are labeled at the highest taxonomy level from genus and ranked by overall contribution to the shared ordination space across the first three principal components. Higher loading magnitudes indicate taxa that most strongly contribute to subject-aware microbial variation captured by CTF.

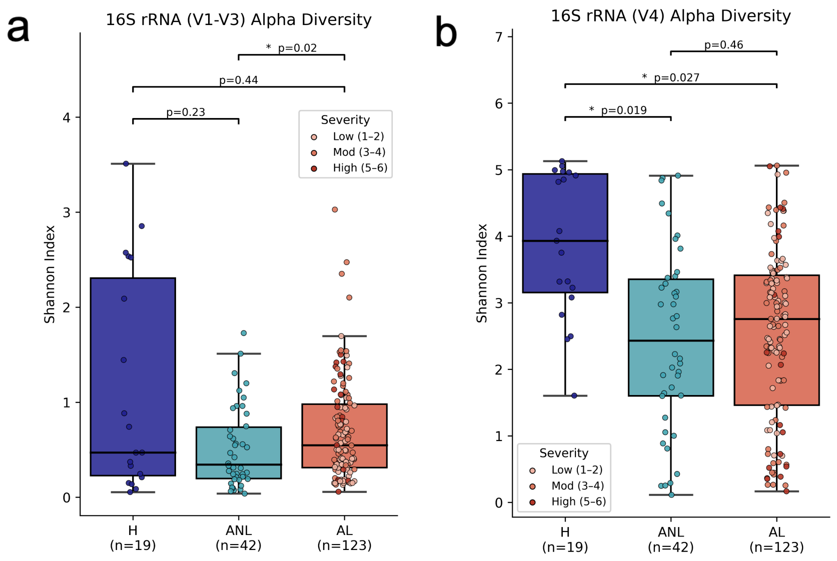


**Suppl. Fig. 3. Alpha and beta diversity analyses between healthy, acne non-lesional, and acne lesional skin groups by Shannon Index. (a–b)** Shannon diversity for **(a)** 16S rRNA V1–V3 and **(b)** 16S rRNA V4. Group differences were assessed using linear mixed-effects models with subject as a random effect; pairwise p-values are shown.

a

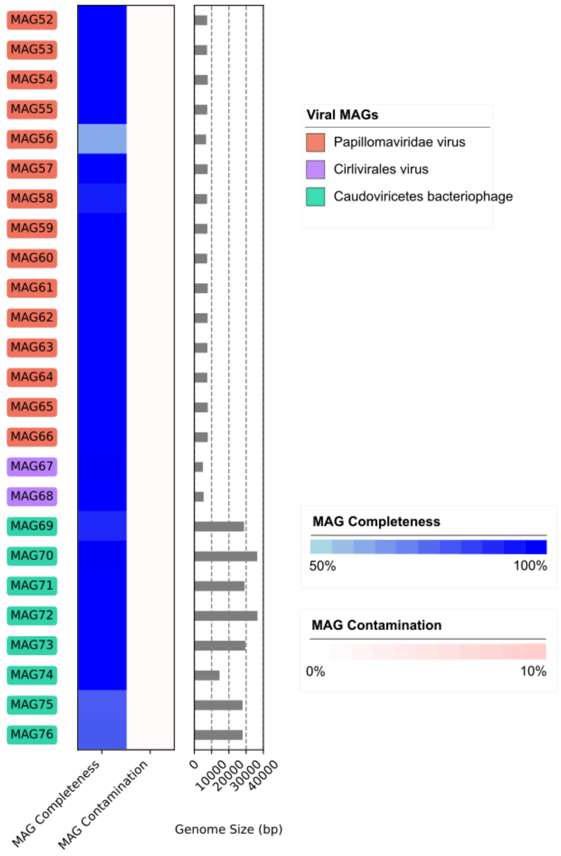


**Suppl. Fig. 4. Viral Metagenome Assembled Genomes identified. (a)** Identification and assessment of 25 viral MAGs belonging to *Papillomaviridae*, *Cirilvirales*, and *Caudoviricetes* (skin bacteriophage).

a

| Gene | Full Name | Description |
|---|---|---|
| aroE | Shikimate 5-dehydrogenase | Involved in the shikimate pathway for aromatic amino acid biosynthesis. |
| atpD | ATP synthase beta chain | Essential for ATP synthesis in cellular respiration. |
| gmk | Guanylate kinase | Catalyzes the phosphorylation of GMP to GDP. |
| guaA | GMP synthase | Plays a role in purine nucleotide biosynthesis. |
| lepA | GTP-binding protein | Involved in protein synthesis and stress response. |
| recA | Recombinase A | Mediates homologous recombination and DNA repair. |
| sodA | Superoxide dismutase | Protects against oxidative damage by scavenging superoxide radicals. |
| tly | Putative hemolysin | Potential virulence factor contributing to hemolytic activity. |
| camp2 | CAMP factor homologue | CAMP factor protein, implicated in virulence and host interaction. |

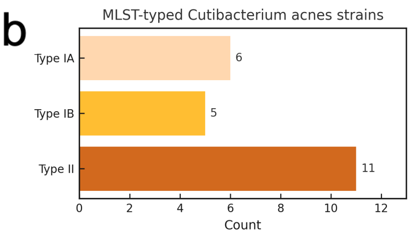


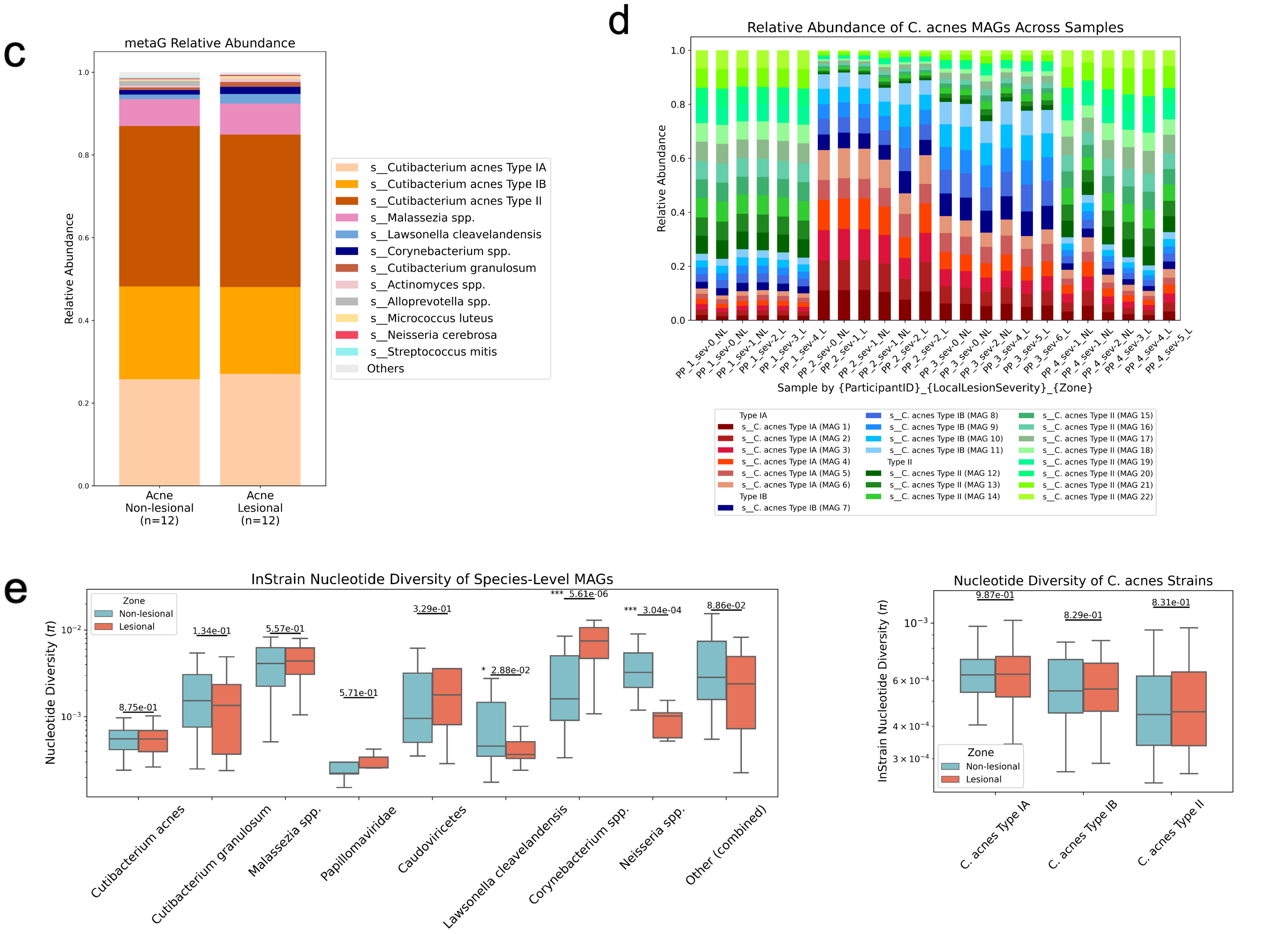


**Suppl. Fig. 5. Strain-level profiling of *C. acnes* and microbial diversity in acne-affected skin. (a)** MLST scheme genes used for *C. acnes* strain classification, including core metabolic, stress response, and virulence loci. **(b)** Distribution of MLST-typed *C. acnes* strains across samples, with Type IA most prevalent. **(c)** Metagenomic abundance of microbial taxa in non-lesional vs. lesional skin (n=12/group), showing *C. acnes* dominance with modest strain-level shifts. **(d)** Per-subject relative abundance of *C. acnes* strains reveals intra- and inter-individual variation, with lesional samples enriched for specific Type IA strains. **(e)** Nucleotide diversity (π) of species-level MAGs shows significantly higher diversity in lesional skin for *Papillomaviridae*, *Caudoviricetes*, *Lawsonella clevelandensis*, and *Corynebacterium*

spp. (Wilcoxon test; $p < 0.05$). **(f)** No significant difference in *C. acnes* strain nucleotide diversity between lesional and non-lesional skin.

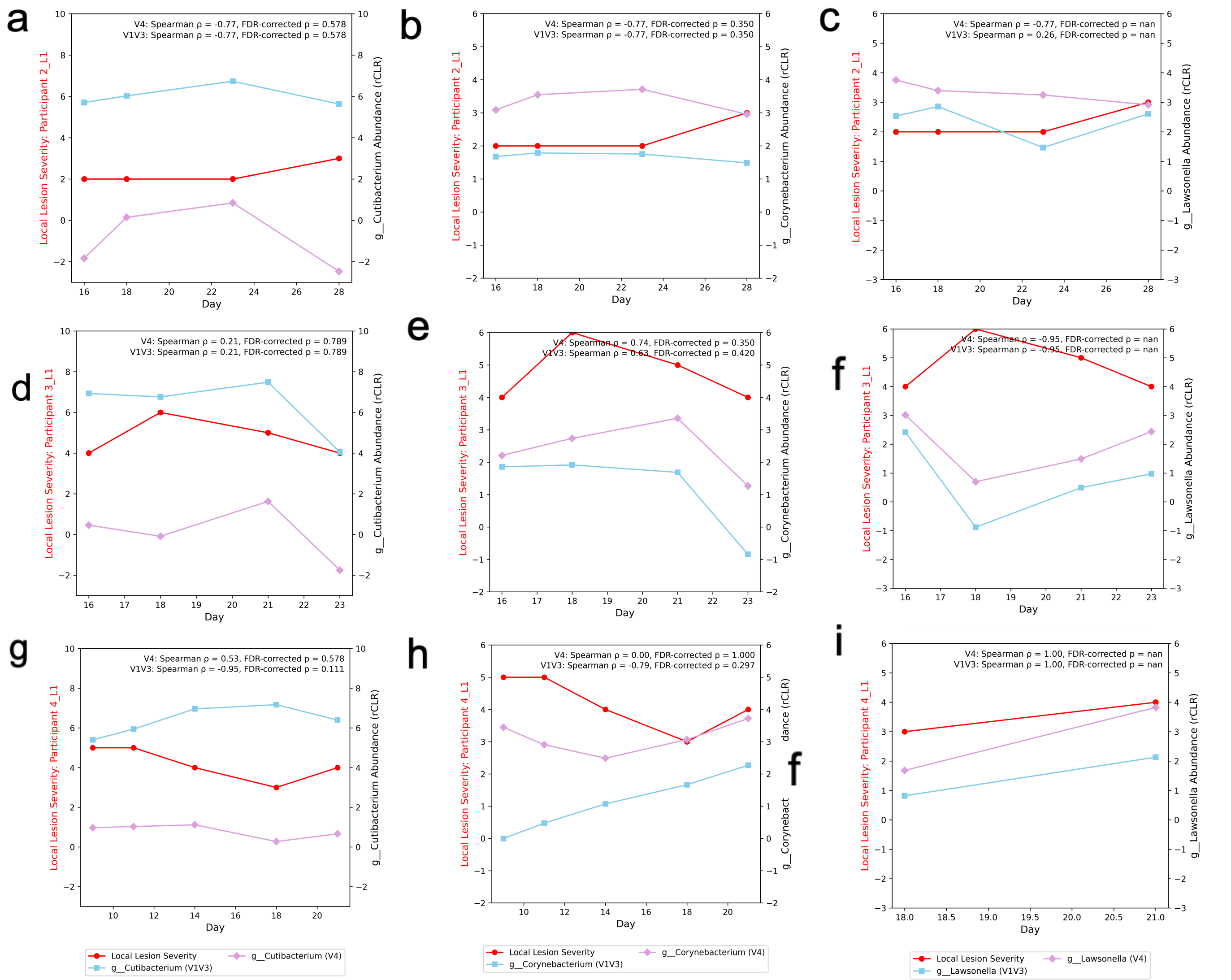


**Suppl. Fig. 6. Longitudinal tracking of acne lesion severity and microbial abundance across individuals highlights subject-specific strain dynamics.** Each panel **(a-i)** shows changes in acne severity (red line, left y-axis) and CLR-transformed abundance of a specific microbe (right y-axis) across time points in an individual subject. Microbial profiles are shown for 16S V1-V3 (light blue), 16S V4 (purple), and shotgun metagenomics (WGS; orange). Panels **(a-c)** display *C. acnes*, *Corynebacterium spp.*, and *Lawsonella* dynamics for Participant 2. Panels **(d-f)** show the same taxa for Participant 3, while panels **(g-i)** depict the same for Participant 4. Spearman correlation coefficients (r) and p-values assess the strength and direction of association between local acne lesion severity and microbial abundance across platforms.

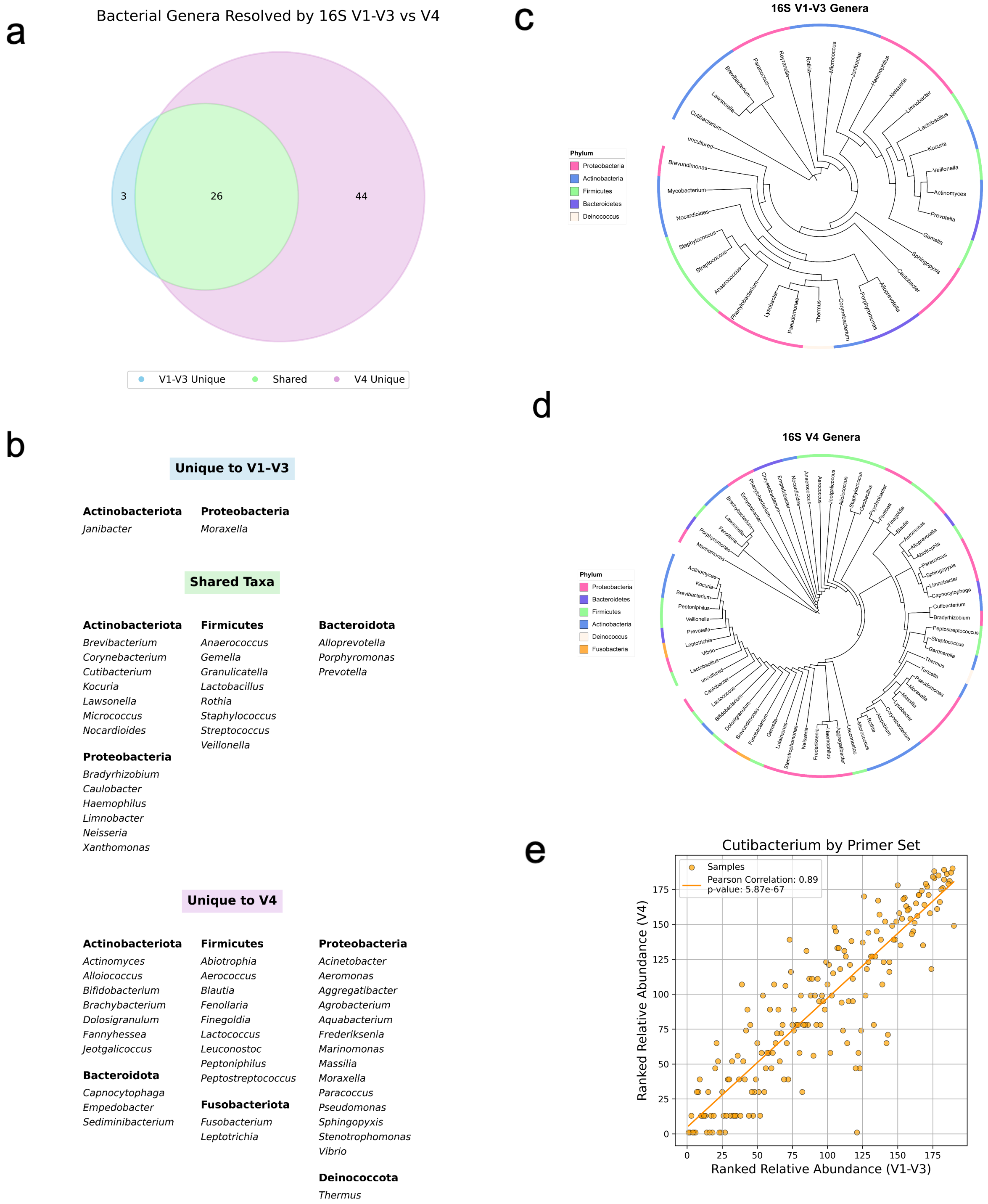
a
Bacterial Genera Resolved by 16S V1-V3 vs V4
3
26
44
V1-V3 Unique
Shared
V4 Unique
b
Unique to V1-V3
Actinobacteriota
Janibacter
Proteobacteria
Moraxella
Shared Taxa
Actinobacteriota
Brevibacterium
Corynebacterium
Cutibacterium
Kocuria
Lawsonella
Micrococcus
Nocardioides
Firmicutes
Anaerococcus
Gemella
Granulicatella
Lactobacillus
Rothia
Staphylococcus
Streptococcus
Veillonella
Bacteroidota
Alloprevotella
Porphyromonas
Prevotella
Proteobacteria
Bradyrhizobium
Caulobacter
Haemophilus
Limnobacter
Neisseria
Xanthomonas
Unique to V4
Actinobacteriota
Actinomyces
Alloiococcus
Bifidobacterium
Brachybacterium
Dolosigranulum
Fannyhessea
Jeotgalicoccus
Bacteroidota
Capnocytophaga
Empedobacter
Sediminibacterium
Firmicutes
Abiotrophia
Aerococcus
Blautia
Fenollaria
Finegoldia
Lactococcus
Leuconostoc
Peptoniphilus
Peptostreptococcus
Fusobacteriota
Fusobacterium
Leptotrichia
Proteobacteria
Acinetobacter
Aeromonas
Aggregatibacter
Agrobacterium
Aquabacterium
Frederiksenia
Marinomonas
Massilia
Moraxella
Paracoccus
Pseudomonas
Sphingopyxis
Stenotrophomonas
Vibrio
Deinococcota
Thermus
c
16S V1-V3 Genera
Phylum
Proteobacteria
Actinobacteria
Firmicutes
Bacteroidetes
Deinococcus
d
16S V4 Genera
Phylum
Proteobacteria
Bacteroidetes
Firmicutes
Actinobacteria
Deinococcus
Fusobacteria
e
Cutibacterium by Primer Set
Samples
Pearson Correlation: 0.89
p-value: 5.87e-67
Ranked Relative Abundance (V4)
Ranked Relative Abundance (V1-V3)
0
25
50
75
100
125
150
175

**Suppl. Fig. 7. Comparison of microbial genera resolved by 16S rRNA V1-V3 vs. V4 primers. (a)** Venn diagram showing genera detected by both primer sets, unique to V1-V3, unique to V4, and their **(b)** taxonomy grouped by phylum. **(c,d)** Maximum likelihood phylogenetic trees of genera identified by V1-V3 **(c)** and V4 **(d)**, with nodes colored by phylum. **(e)** Concordance of *Cutibacterium* relative abundance rankings between 16S rRNA gene sequencing using V1-V3 and V4 primer sets (Pearson correlation $r = 0.89$, $p = 5.87e{-}67$).

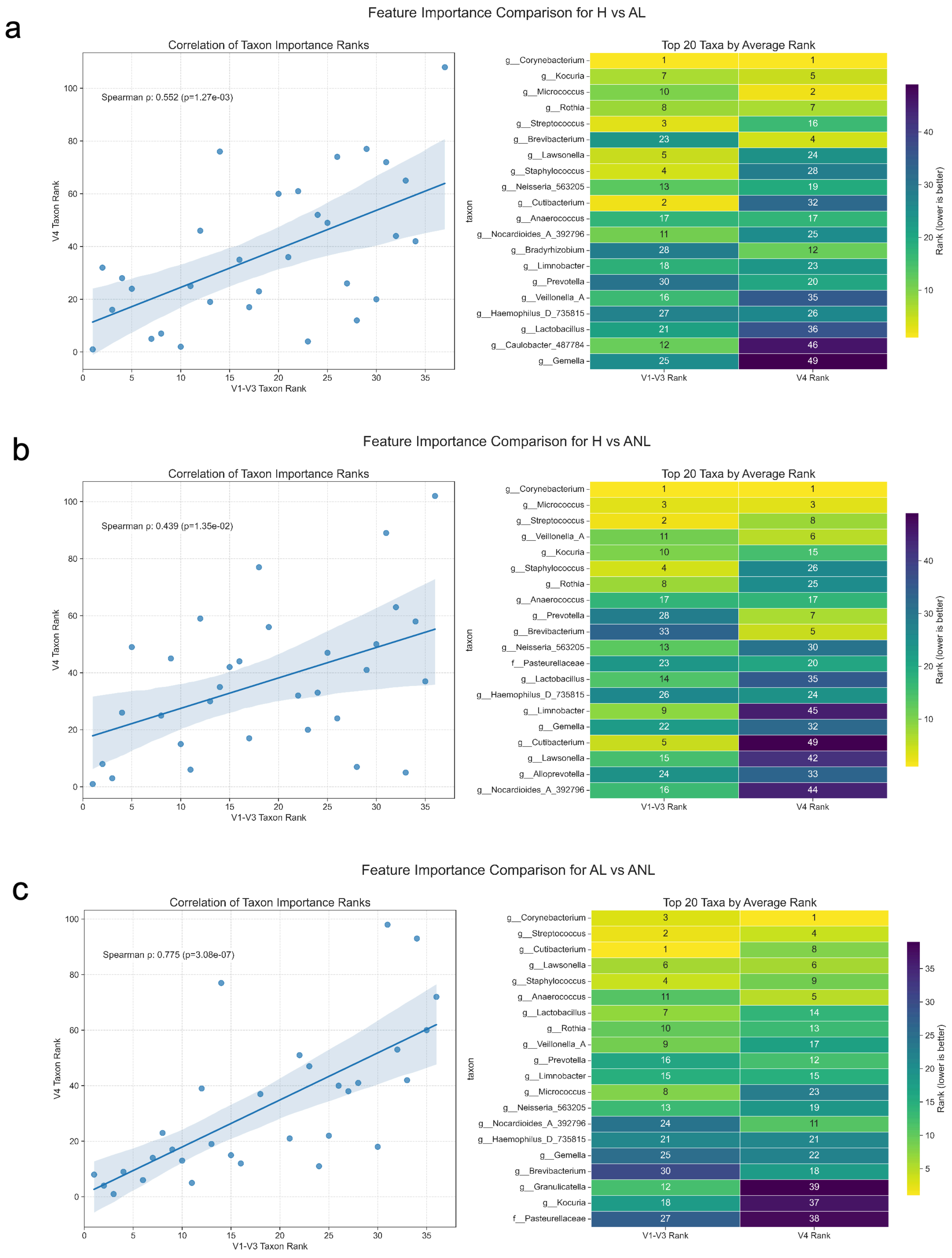
a
Feature Importance Comparison for H vs AL
Correlation of Taxon Importance Ranks
Spearman ρ: 0.552 (p=1.27e-03)
V4 Taxon Rank
V1-V3 Taxon Rank
Top 20 Taxa by Average Rank
taxon
V1-V3 Rank
V4 Rank
Rank (lower is better)
g__Corynebacterium 1 1
g__Kocuria 7 5
g__Micrococcus 10 2
g__Rothia 8 7
g__Streptococcus 3 16
g__Brevibacterium 23 4
g__Lawsonella 5 24
g__Staphylococcus 4 28
g__Neisseria_563205 13 19
g__Cutibacterium 2 32
g__Anaerococcus 17 17
g__Nocardioides_A_392796 11 25
g__Bradyrhizobium 28 12
g__Limnobacter 18 23
g__Prevotella 30 20
g__Veillonella_A 16 35
g__Haemophilus_D_735815 27 26
g__Lactobacillus 21 36
g__Caulobacter_487784 12 46
g__Gemella 25 49
b
Feature Importance Comparison for H vs ANL
Correlation of Taxon Importance Ranks
Spearman ρ: 0.439 (p=1.35e-02)
V4 Taxon Rank
V1-V3 Taxon Rank
Top 20 Taxa by Average Rank
taxon
V1-V3 Rank
V4 Rank
Rank (lower is better)
g__Corynebacterium 1 1
g__Micrococcus 3 3
g__Streptococcus 2 8
g__Veillonella_A 11 6
g__Kocuria 10 15
g__Staphylococcus 4 26
g__Rothia 8 25
g__Anaerococcus 17 17
g__Prevotella 28 7
g__Brevibacterium 33 5
g__Neisseria_563205 13 30
f__Pasteurellaceae 23 20
g__Lactobacillus 14 35
g__Haemophilus_D_735815 26 24
g__Limnobacter 9 45
g__Gemella 22 32
g__Cutibacterium 5 49
g__Lawsonella 15 42
g__Alloprevotella 24 33
g__Nocardioides_A_392796 16 44
c
Feature Importance Comparison for AL vs ANL
Correlation of Taxon Importance Ranks
Spearman ρ: 0.775 (p=3.08e-07)
V4 Taxon Rank
V1-V3 Taxon Rank
Top 20 Taxa by Average Rank
taxon
V1-V3 Rank
V4 Rank
Rank (lower is better)
g__Corynebacterium 3 1
g__Streptococcus 2 4
g__Cutibacterium 1 8
g__Lawsonella 6 6
g__Staphylococcus 4 9
g__Anaerococcus 11 5
g__Lactobacillus 7 14
g__Rothia 10 13
g__Veillonella_A 9 17
g__Prevotella 16 12
g__Limnobacter 15 15
g__Micrococcus 8 23
g__Neisseria_563205 13 19
g__Nocardioides_A_392796 24 11
g__Haemophilus_D_735815 21 21
g__Gemella 25 22
g__Brevibacterium 30 18
g__Granulicatella 12 39
g__Kocuria 18 37
f__Pasteurellaceae 27 38

**Suppl. Fig. 8. Comparison of microbial feature importance rankings across 16S V1-V3 and V4 datasets in acne phenotype classification. (a)** Feature importance comparison for distinguishing H vs. ANL samples. **(b)** Feature importance comparison for AL vs. ANL. **(c)** Feature importance comparison for AL vs. H. Left panels show the correlation of feature importance ranks for microbial taxa identified by machine learning classifiers trained on V1-V3 and V4 16S rRNA datasets. Each point represents a shared feature (taxon), with Spearman correlation coefficients reported. Trendlines and 95% confidence intervals are shown in red. The right panels show the top 20 features ranked by average importance across both primer sets. Features are annotated at the family (*f*__) and genus (*g*__) levels. Heatmaps display feature ranks from V1-V3 and V4 datasets, with darker shades indicating higher importance (lower numerical rank).

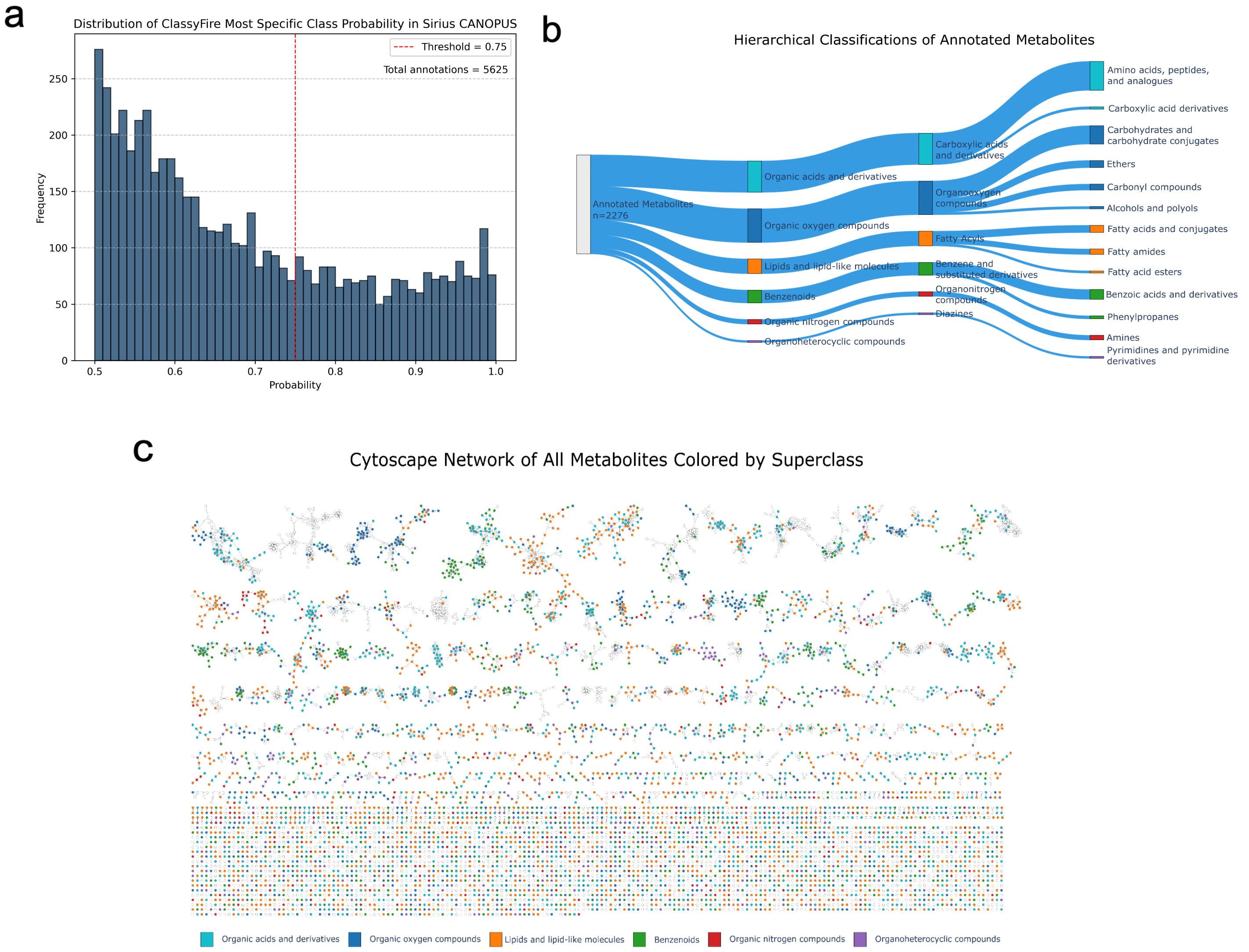


**Suppl. Fig. 9. Metabolomic annotation and chemical classification. (a)** Distribution of SIRIUS CANOPUS annotation probabilities for the most specific ClassyFire chemical class (n=5,625 annotations). The red dashed line indicates the 75% confidence threshold for high-quality annotations. **(b)** Sankey diagram showing hierarchical ClassyFire classification (Superclass, Class, Subclass) of high-confidence metabolites (probability ≥75%; n = 2,276). **(c)** Molecular network of 2,276 metabolites organized by structural similarity (shared MS/MS fragmentation patterns) and colored by ClassyFire superclass: organic acids and derivatives (light blue), organic oxygen compounds (dark blue), lipids and lipid-like molecules (orange), benzenoids (green), organic nitrogen compounds (red), organoheterocyclic compounds (purple). Nodes represent metabolites; edges indicate structural similarity.

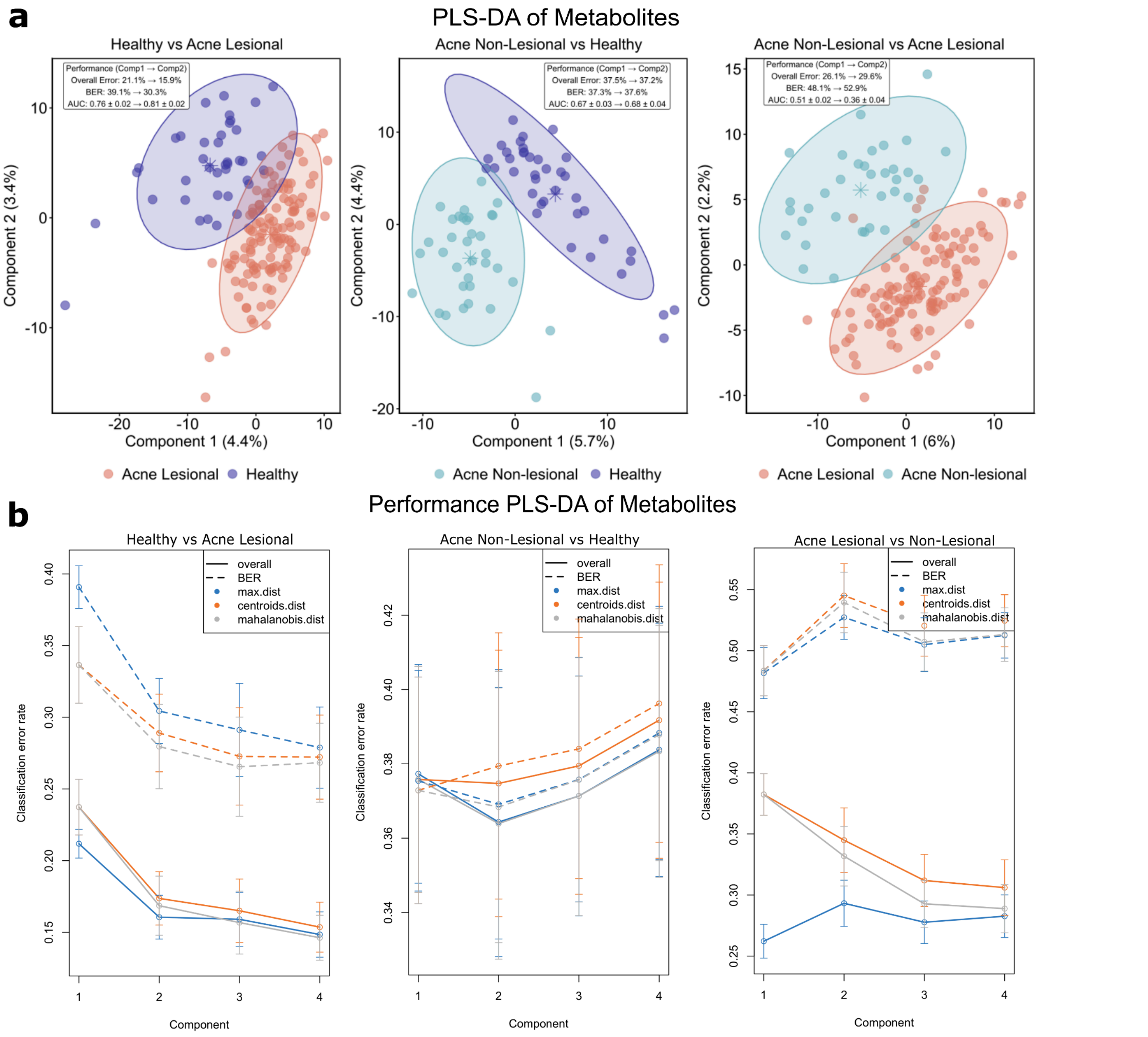


**Suppl. Fig. 10. PLS-DA classification of acne phenotypes using metabolomic profiles. (a)** PLS-DA ordination plots for pairwise comparisons: H vs. AL (left), ANL vs. H (center), and AL vs. ANL (right). Ellipses represent 95% confidence intervals. Performance metrics: overall error rate, balanced error rate (BER), and area under the curve (AUC). **(b)** PLS-DA classification performance across 1-4 components, evaluated by overall error, BER, maximum distance, centroid distance, and Mahalanobis distance. Error bars show standard deviation across cross-validation folds. H vs. AL achieved best performance, with error rates improving as components increased, demonstrating that discriminative metabolites effectively separate acne phenotypes.

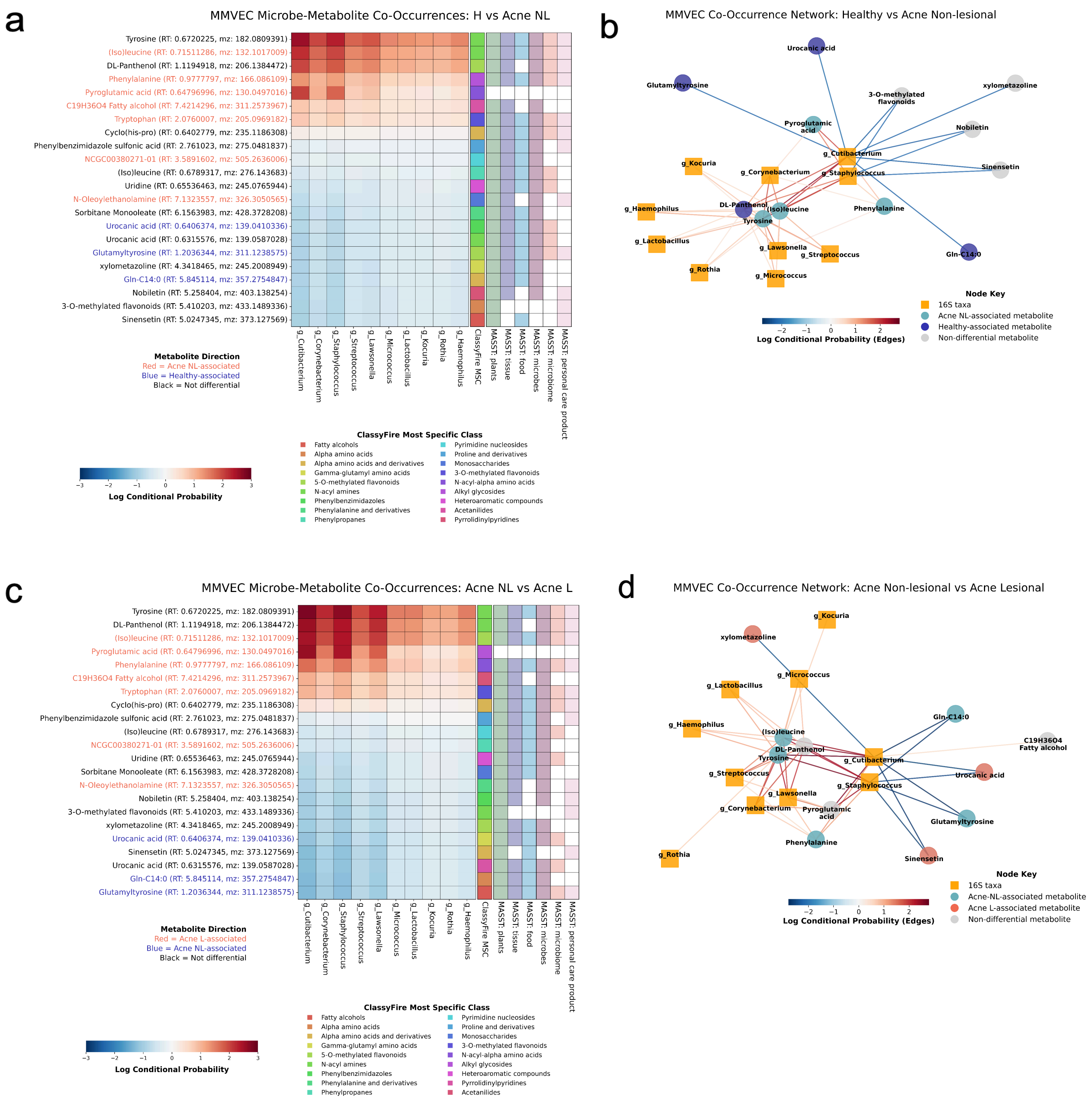


**Suppl. Fig. 11. Microbe-metabolite co-occurrences across acne skin phenotypes. (a,b)** Healthy vs. acne non-lesional: **(a)** Heatmap of log conditional probabilities between top 30 metabolites and 10 key taxa. Color coding: red=acne non-lesional, blue=healthy, black=non-differential. **(b)** Network of top 20% associations; taxa (orange squares) and metabolites (circles) colored by VIP association. **(c,d)** Acne non-lesional vs. lesional: heatmap and network as in **(a,b)**, comparing non-lesional (blue) vs. lesional (red).

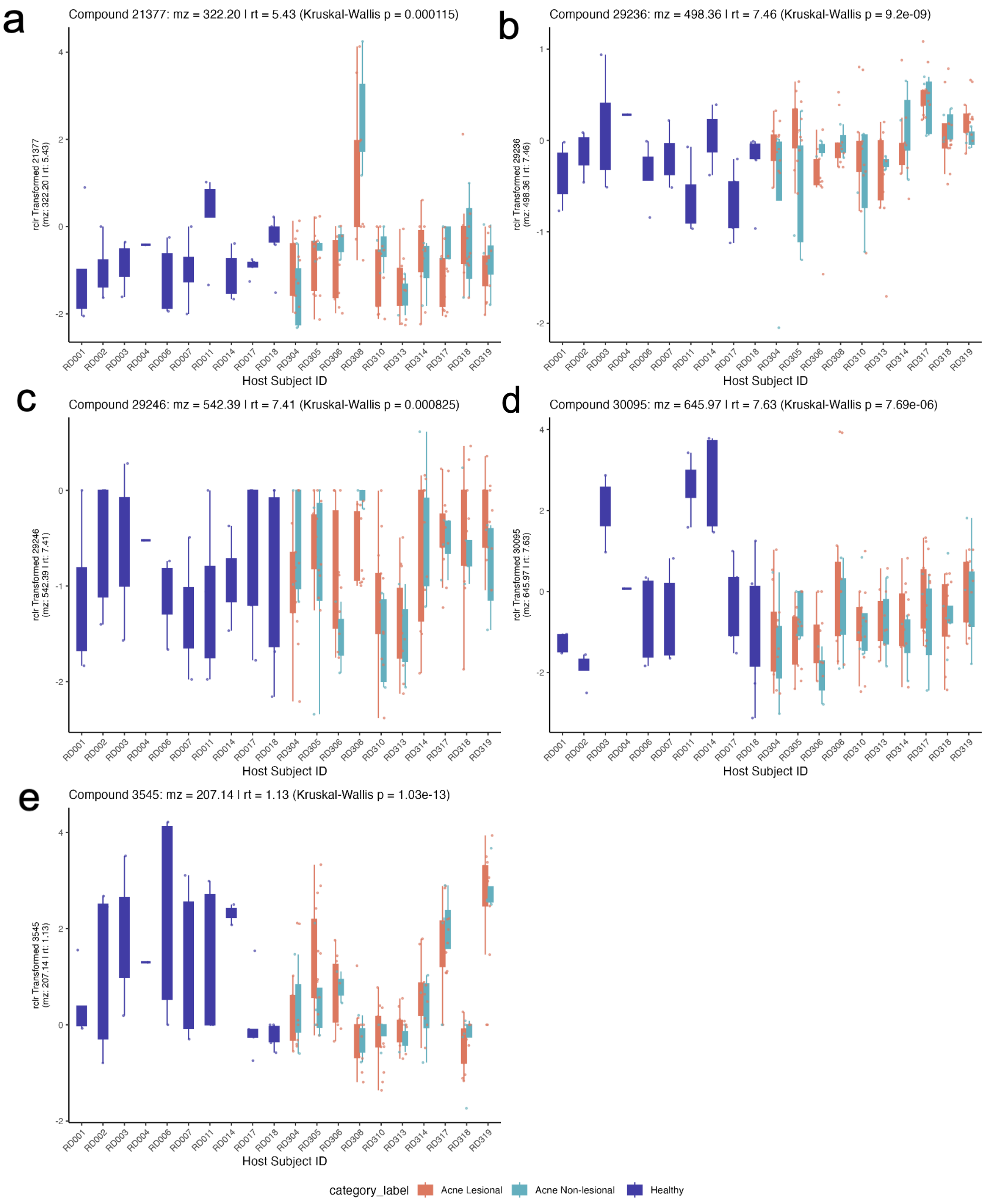


**Suppl. Fig. 12. RCLR-transformed abundances of significant metabolites matched to the personal care products database, identified through PLS-DA analysis.** Abundances are shown per host subject ID for **(a)** Compound 21377, **(b)** Compound 29236, **(c)** Compound 29246, **(d)** Compound 30095, and **(e)** Compound 3545. The mass-to-charge ratio (m/z) and retention time (RT) are indicated for each compound. Statistical significance was assessed using the Kruskal-Wallis test.

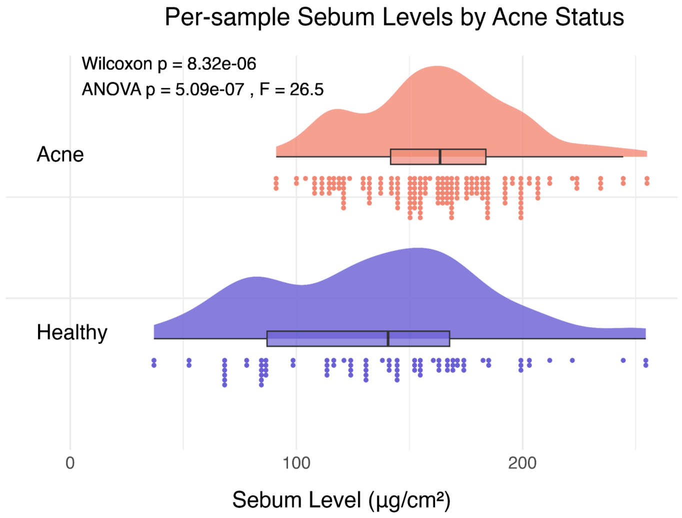


**Suppl. Fig. 13. Distribution of sebum levels by acne status.** Sebum measurements (µg/cm²) for acne (red) and healthy (blue) groups are shown as individual data points with overlaid density plots and box plots (median and interquartile range). Both Wilcoxon (*p = 8.32e-06*) and ANOVA (*p = 5.09e-07, F = 26.5*) tests indicate significantly higher sebum levels in the acne group.
